# Cofilin-Driven Nuclear Deformation Drives Dendritic Cell Migration through the Extracellular Matrix

**DOI:** 10.1101/2023.07.10.548429

**Authors:** Harry Warner, Giulia Franciosa, Guus van der Borg, Felix Faas, Claire Koenig, Rinse de Boer, René Classens, Sjors Maassen, Maksim Baranov, Shweta Mahajan, Deepti Dabral, Britt Coenen, Frans Bianchi, Niek van Hilten, Herre Jelger Risselada, Wouter H. Roos, Jesper Olsen, Laia Querol Cano, Geert van den Bogaart

## Abstract

To mount an adaptive immune response, dendritic cells must process antigens, migrate to lymph nodes and form synapses with T cells. Critical to 3D migration and mechano-sensing is the nucleus, which is the size-limiting barrier for navigation through gaps in the extracellular matrix. Here, we show that inflammatory activation of dendritic cells leads to the nucleus becoming spherically deformed, adopting a raison-like shape and enables dendritic cells to overcome the typical 2 – 3-micron pore limit for 3D-migration. We show that the nuclear shape-change is partially attained through reduced cell adhesion, whereas improved migration through extracellular matrix is achieved through reprogramming of the actin cytoskeleton. Specifically we show that cofilin-1 is phosphorylated at serine 41 drives the assembly of a Cofilin-ActoMyosin (CAM)-ring proximal to the nucleus and enhancing migration through 3D collagen gels. In summary, these data describe novel signaling events through which dendritic cells simultaneously deform their nucleus and enhance their migratory capacity; molecular events that may be re-capitulated in other contexts such as wound healing and cancer.

## Introduction

Dendritic cells form the interface between the innate and adaptive immune systems and operate throughout the body in environments with very different stiffnesses. Before activation, dendritic cells reside in virtually all tissues within the body where they adapt to the local biochemical and physical cues provided by surrounding cells and the extracellular matrix (ECM)(Patente et al., 2019; Park *et al*., 2021; Chakraborty *et al*., 2021; Meddens *et al*., 2016). Therefore, dendritic cells must be extremely plastic in their mechanosensing behavior; being able to function at stiffnesses ranging from <1 kPa (e.g., brain) to >10 kPa (e.g., muscle). Moreover, following the recognition and ingestion of a pathogen, dendritic cells become activated and rapidly migrate to a local lymph node to activate T cells. To facilitate this migration and adapt to the physiological properties of lymph nodes, activated dendritic cells undergo both rapid and systematic re-programming of their adhesion machinery, breaking down podosomes and becoming less adhesive(West *et al*., 2008). Furthermore, once at a lymph node, dendritic cells stimulate fibroblasts to trigger lymph node expansion (Acton *et al*., 2014; Martinez *et al*., 2019). Lymph node expansion drives stiffening of the lymph node, and prevents crowding of lymphocytes within the lymph node; a mechanical event that dendritic cells have to rapidly adapt to as well (Assen *et al*., 2021).

Cell-matrix adhesion is critical to most metazoan life(Hynes, 2012). In particular modifications to cell-matrix adhesion efficiency via cell spreading are known to be far reaching, regulating not only cell shape, but gene expression and cell-differentiation (McBeath *et al*., 2004; Connelly *et al*., 2010; Murray *et al*., 2013; Carley *et al*., 2021; de Belly *et al*., 2021). Furthermore, forces from the ECM are known to be transmitted by the actin cytoskeleton to the nucleus via LINC (linker of nucleoskeleton and cytoskeleton) complexes. (Maniotis, Chen and Ingber, 1997; Khatau *et al*., 2009; Lovett *et al*., 2013; Vishavkarma *et al*., 2014; Chen, Co and Ho, 2015). This can regulate nuclear morphology (with the nucleus getting more round as adhesion reduces) and has even been reported to regulate cell differentiation (Carley et al., 2021). Furthermore, multiple studies have indicated that dendritic cell migration can occur in the absence of integrin-based adhesion machinery (Lämmermann *et al*., 2008a; Reversat *et al*., 2020), and numerous results have indicated that activated dendritic cells indeed switch to a less adhesive migratory phenotype(Van *et al*., 2006; West *et al*., 2008). Therefore, it seems possible that reduced force transmission from the extracellular matrix to the dendritic cell nucleus may occur following inflammatory activation.

Critical to both 3D migration and mechano-sensing is the nucleus. During migration, the nucleus is used by cells to probe gap-sizes in the ECM, transmit signals of confinement to the actin cytoskeleton and generate intracellular pressure (Renkawitz *et al*., 2019) (Lomakin *et al*., 2020) (Venturini *et al*., 2020) (Petrie, Koo and Yamada, 2014). Nesprin-2G based LINC complexes connect the actin cytoskeleton, via Nesprin-2G and SUN proteins, to lamin filaments, enabling the cell to pull the nucleus through gaps in the ECM and mechanoregulate the genome (Davidson *et al*., 2020; Gilbert *et al*., 2019; Nava *et al*., 2020). Furthermore, the nucleus is known to act as a size limiting organelle, blocking migration of mononuclear cells (including dendritic cells) through ECM gaps smaller than about 2 – 3 microns (Wolf *et al*., 2013; Baranov *et al*., 2014) (Thiam *et al*., 2016). Whilst the role of the nucleus in dendritic cell migration is well defined, how the dendritic cell nucleus senses force when tissue-resident is poorly understood, as is the effect of inflammatory activation on the nucleus in the context of 3D-migration.

Here, using quantitative microscopy we demonstrate that dendritic cell activation leads to a change in nuclear morphology and enhanced passage of the nucleus (and thus the whole cell) through 2-micron pores. Whilst activation-induced de-adhesion can explain the change in dendritic cell nuclear morphology it cannot explain the increased migration efficiency, as atomic force microscopy (AFM) measurements indicate the bulk nucleus stiffens in response to inflammatory stimulation. Instead, we demonstrate using a combination of phosphoproteomics and quantitative microscopy that cofilin-1 is phosphorylated at serine 41 in response to inflammatory activation and that this drives the assembly of a contractile cofilin-actomyosin (CAM)-ring around the nucleus to facilitate enhanced nuclear squeezing and increased migration efficiency through the ECM.

## Results

As inflammatory activation has long been known to reduce dendritic cell adhesion in response to LPS, it is logical that activation may concomitantly alter the shape of the dendritic cell nucleus. Therefore, to examine how inflammation-induced de-adhesion influences the dendritic cell nucleus, we differentiated monocyte-derived dendritic cells (moDCs) from human peripheral blood monocytes, stimulated them with bacteria-derived lipopolysaccharide (LPS) overnight, and quantified the *Z*-projected area of complete dendritic cells as well as dendritic cell nuclei. (**Fig. 1A**). This revealed that, in the *X-Y* plane, dendritic cells become less spread (consistent with previous results (Van *et al*., 2006; West *et al*., 2008)) and the *Z-*projected areas of the nuclei become smaller in response to LPS stimulation (**Fig. 1B**). Through the use of 3D airyscan super-resolution microscopy, we measured the approximate volume of nuclei from inactivated and activated dendritic cells (**Fig. 1C and 1D**). This revealed that although the nuclei get smaller in the projected *X*-*Y* plane upon activation, they maintain a constant volume; they extend into the *Z*-plane (herein referred to as “nuclear deformation”).

**Figure 1.**
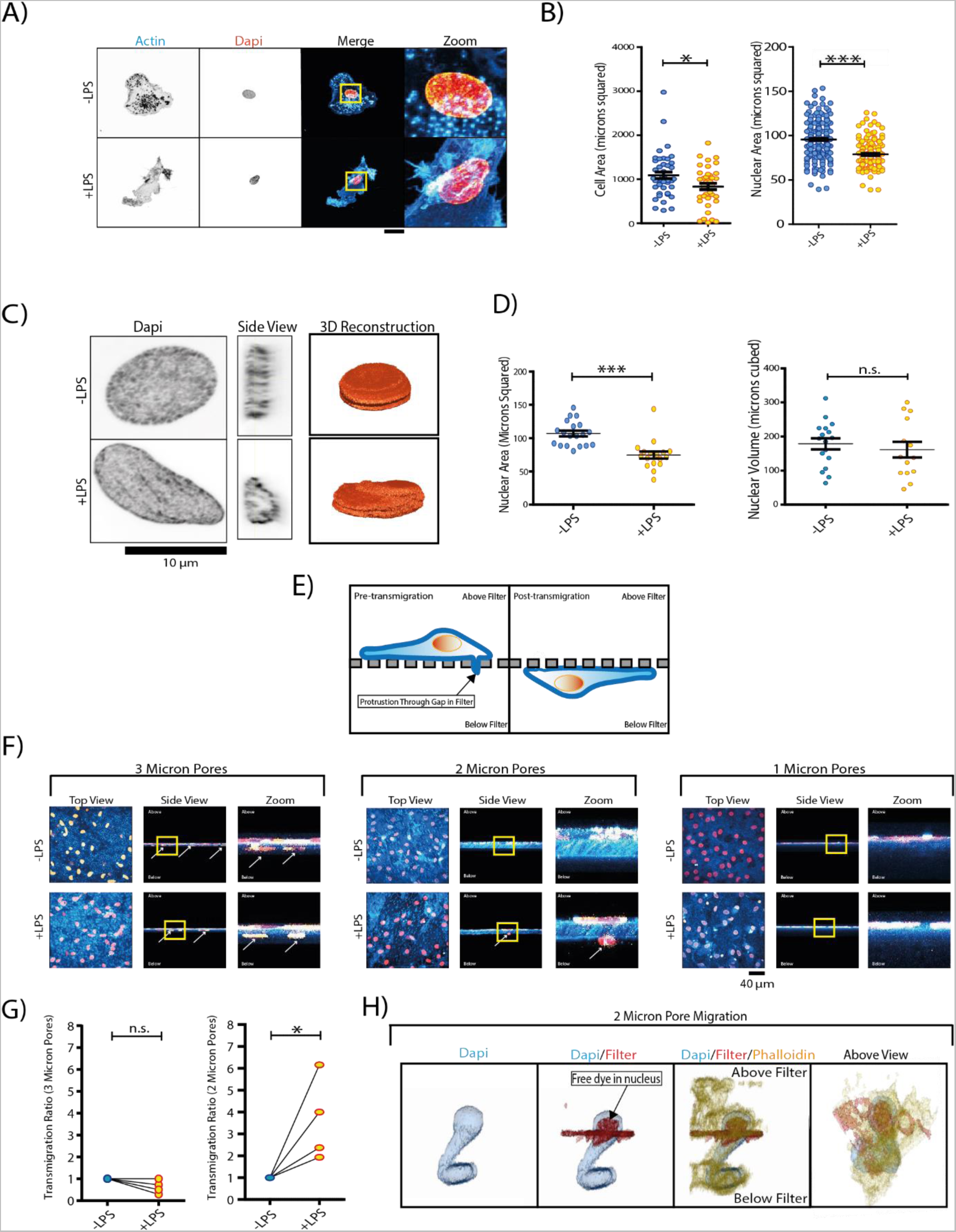
LPS-Stimulation Spherically Deforms the Dendritic Cell Nucleus and Enhances Transmigration. A) Confocal micrographs of dendritic cells stained with phalloidin (cyan in merge) and DAPI (orange) and cultured overnight in the presence or absence of LPS. Activated dendritic cells present a deformed shriveled nucleus in comparison to inactive dendritic cells. B) Quantification of Z-projected cellular area (cell spreading) (minimum 39 measurements per condition, 3 biological repeats) and nuclear area (minimum 157 measurements per condition, 3 biological repeats). C) Airyscan and 3D reconstruction of dendritic cell nuclei following overnight culturing in the presence or absence of LPS. Activated dendritic cell nuclei appear taller. D) Quantification of Z-projected nuclear size and quantification of nuclear volume (of the same nuclei) reveals a change in X-Y area, but not volume (at least 14 measurements per condition over 4 biological repeats). E) Schematic demonstrating filter experiment setup. Dendritic cells are seeded on top of gelatin-coated polycarbonate filters with either 3, 2 or 1 µm pores, through which they can protrude and potentially transmigrate. F) Confocal micrographs of dendritic cells cultured on gelatin-coated polycarbonate filters with set pore sizes (fluorescent gelatin in blue, dapi in red). G) Quantification of the normalized transmigration ratio of dendritic cells through 3-micron pores and 2-micron pores, over a 12 hour + period, in the presence or absence of LPS (defined as the number of cells passing through the filter, normalised to local seeding density). Data for 4 donors. H) 3D reconstruction of a dendritic cell observed migrating through a 2-micron filter, in the presence of LPS. Data information: Scale bars indicate 20 microns. Statistical significance was calculated using 2-sided unpaired t-tests (selected according to distribution pattern of the data). *, P < 0.05; ***, P < 0.001; n.s., not significant. Error Bars= standard error mean (SEM). For Box and Whisker plots, Box represents 25^th^ to 75^th^ percentiles, whiskers represents maximum and minumum values, middle band represents data median,+ represents data mean.

Having established that the dendritic nucleus deforms in response to LPS, we also examined the effect of LPS stimulation on CD14^+^ monocyte-derived macrophages (**Fig. S1A and S1B**). We compared macrophages with dendritic cells, because, in contrast to dendritic cells, the activation of macrophages by inflammatory stimuli does not result in their migration to the lymph nodes. Opposite to our findings with dendritic cells, the *Z*-projected area of the nuclei of macrophages became larger, suggesting that the nuclear morphology change observed in dendritic cells is required for a dendritic cell-specific function. We also verified that the same nuclear deformation occurs in dendritic cells in response to bacterial exposure; a scenario in which dendritic cells are exposed to multiple inflammatory ligands. Exposure of dendritic cells to intact *Escherichia coli* bacteria produced a similar shift in nuclear morphology (**Fig. S1C and S1D**).

We also confirmed that macrophages did not contaminate our moDC cultures: monocytes purified from primary donors are relatively uncontaminated by cells expressing macrophage markers (which overlap with moDCs) (**Fig. S1E – S1H**), suggesting that macrophage contamination is very low in our initial, pre-differentiated cultures. Furthermore, stimulation of dendritic cells with LPS did not largely affect seeding density (**Fig SI**), meaning that activation did not largely bias the results to more adherent contaminant cells. Thus macrophage contaminations are unlikely to impact our results.

We hypothesized that activation-associated changes to the nucleus likely facilitate more efficient migration through gaps in the ECM, as dendritic cells have to migrate to lymph nodes to perform their function, whereas macrophages do not. We first confirmed that activation increases dendritic cell migration efficiency in 3D matrix. In order to do this we seeded inactive and active dendritic cells into bovine dermal collagen produced at a concentration of 1.7 mg.ml^−1^ and analysed their mean square displacement (MSD). We selected MSD as this is a principal means to measure the confinement of a particle (i.e. a dendritic cell) within a closed environment (i.e. the collagen gel). (**Fig. S1J – S1O**). This confirmed that inflammatory activation enhances migration in 3D collagen, as the MSD significantly increased in activated dendritic cells compared to inactive dendritic cells (**Fig. S1O**). Furthermore, previous work has demonstrated that 3D matrix made of bovine dermal collagen at 1.7mg ml^−1^ has pore sizes ranging from ~2-6 microns (i.e. approaching the size limit for 3D migration as determined by the nucleus) (Wolf *et al*., 2013). Thus this result implies that re-programming of the nucleus enhances the increased migration capacity of activated dendritic cells.

To test the contribution of the nucleus more directly to this increased migration efficiency, we utilized an approach previously used to study the impact of the nucleus on migration through gaps in the ECM. Specifically, dendritic cells were seeded onto polycarbonate filters coated with fluorescent gelatin, with defined pore sizes of 3, 2 and 1 microns, in the absence and presence of LPS (**Fig. 1E**). These pore sizes were selected as the nucleus typically limits migration through ECM gaps that are 3 microns or larger (Wolf *et al*., 2013; Baranov *et al*., 2014). Furthermore, this experimental setup enabled us to partially mimic the ECM, whilst simultaneously blocking the cell’s ability to manipulate the size of pores in the ECM (as the polycarbonate filters cannot themselves be physically altered by the cells). Although this setup does not represent a scenario likely found *in vivo*, it does allow us to directly test the impact of the nucleus on migration through gaps in the ECM. Consistent with previous work (Baranov *et al*., 2014; Thiam et al., 2016), in the absence of LPS, dendritic cells were able to migrate through 3-micron pores, but were largely (but not completely) blocked from migrating through 2-micron pores and completely blocked from migrating through 1-micron pores. However, in the presence of LPS dendritic cells were able to migrate with far greater efficiency through 2-micron pores, but were still blocked from migrating through 1-micron pores **(Fig. 1F – 3H)**. We conclude that the activation process enables more efficient passage of the dendritic cell nucleus through gaps in the ECM, and thus activation-induced changes to the nucleus are not purely coincidental. Therefore it is important to understand the molecular mechanisms that re-program the dendritic cell nucleus during inflammatory activation; these may be required for activation of the adaptive immune system and redeployed by other cell types such as metastatic cancer cells.

We also verified that changes to the nucleus in fibronectin-based gel are not due to the biochemical input of ECM by seeding dendritic cells on glass supports coated with various ECM components (collagen, fibronectin, laminin, vitronectin, or fibrinogen), with and without LPS (**Fig. S2A - S2J**). This indicated that in 2D culture, the biochemical input of most of these ECM components does neither impact dendritic cell nuclear morphology nor the LPS induced spherical deformation. Laminin coating however overrode LPS-induced nuclear deformation. This suggests that *in vivo* basement membranes transmit biochemical signals to the nuclear envelope.

To verify that nuclear deformation is at least partially due to reduced dendritic cell adhesion and to confirm that this change can occur at physiological stiffnesses, we examined the morphology of dendritic cell nuclei of cells, with and without LPS, cultured on fibronectin-gels with set stiffness values. These stiffnesses ranged from 10 kPa, through 5 kPa to 1 kPa. These values were selected to broadly reflect the stiffness of tissues ranging from muscle to liver respectively. The *Z-*projected nuclear areas of the inactive dendritic cells continued to get smaller (resembling activated nuclei in dendritic cells cultured on glass) as the substrate became softer until they reached the 1 kPa gels, at which point they became larger (**Fig. 2A and 2B**). Activation with LPS reduced the *Z*-projected area of the nuclei, relative to inactive dendritic cells, on both glass and 10 kPa matrix, but failed to show a difference on 5 kPa and softer matrices. These results imply that the deformation of the nucleus in response to LPS is due to reduced dendritic cell adhesion, and demonstrate that the difference between inactive and active dendritic cell nuclei can be observed within a physiological stiffness range (**Fig. 2A and 2B**). As LPS stimulation-induced nuclear deformation was dependent on alterations to the cell’s adhesive properties, we hypothesized that activation may (at least in part) alter mechanical forces across actin-based LINC complexes. To examine this, we used the Nesprin-2 tension probe mN2G-TS (Arsenovic and Conway, 2018). This probe has an internal mTFP-Venus donor/acceptor Förster resonance energy transfer (FRET) pair, which is pulled apart as mechanical tension across the protein increases (**Fig. 2C**) (Arsenovic and Conway, 2018). Thus, the FRET efficiency increases as tension across the Nesprin-2G-based LINC complexes decreases. The mN2G-TS probe revealed a reduction in mechanical tension across nesprin-2 based LINC complexes in response to LPS **(Fig. 2D and 2E)**. We also verified the accuracy of the probe by comparing the FRET ratio produced by the WT probe, compared to the headless Nesprin-2 tension probe that is unable to connect to the actin cytoskeleton resulting in maximum FRET (Arsenovic and Conway, 2018) (**Fig. S3A and S3B**). We verified these results using airsycan microcopy of F-actin. This revealed the presence of an actin cap, as has been observed in multiple eukaryotic cells, including dendritic cells (Woroniuk *et al*., 2018; Gaertner *et al*., 2022), and may be utilised by inactive dendritic cells to squeeze through gaps in the ECM (Thiam *et al*., 2016). This cap is extremely fine, and is difficult to observe through conventional confocal microscopy. We also examined the intermediate filament vimentin, as this has been shown to support the nucleus in dendritic cells and is supported by the F-actin cytoskeleton (Sutoh Yoneyama *et al*., 2014)(Da *et al*., 2020)(Duarte *et al*., 2019)(Serres *et al*., 2020). Both actin and vimentin cytoskeletons showed a reduced intensity above the nucleus in response to LPS (**Fig. 2F – 2I**). We also examined the microtubule cytoskeleton (**Fig. S3C and S3D**). We observed a slight, but non-significant reduction in the association of microtubules with the nucleus, in response to LPS.

**Figure 2.**
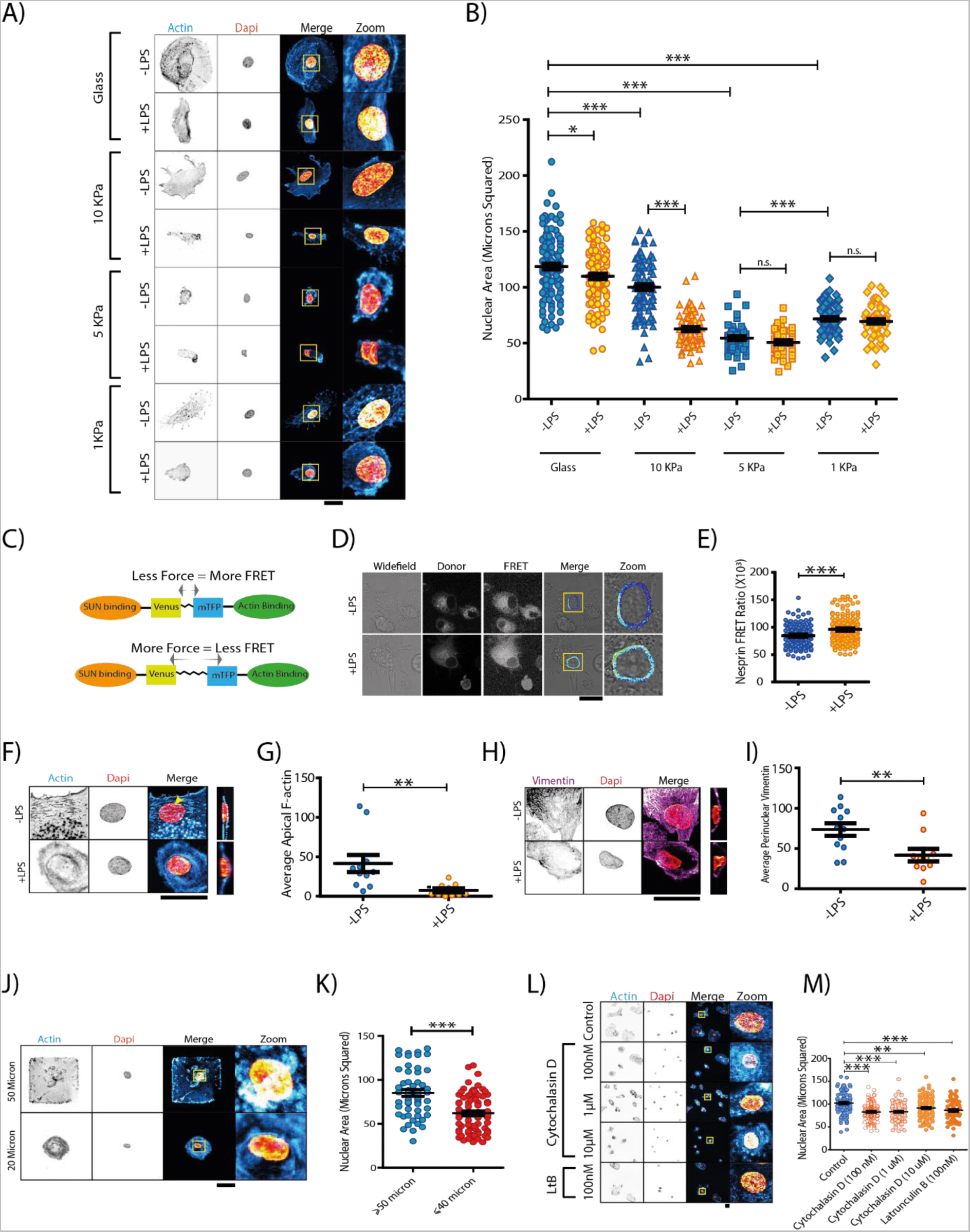
LPS-Stimulation Drives Nuclear Deformation through Adhesion Loss. A) Confocal micrographs of dendritic cells stained with phalloidin (cyan in merge) and DAPI (orange) and cultured on substrates of various stiffnesses in the presence or absence of LPS (overnight). These stiffnesses broadly reflect tissues ranging from muscle to liver. B) Quantification of the Z-projected nuclear areas in the absence or presence of LPS, in dendritic cells cultured on substrates of various stiffnesses (minimum of 44 measurements per condition over 4 biological repeats). C) Schematic of the Nesprin-2 FRET sensor. D) Example ratiometric FRET image of Nesprin-2 probe in the presence or absence of LPS (3 hours plus). E) Quantification of FRET Ratio at the dendritic cell nuclear membrane. (156 measurements across 4 donors) F) Airyscan images of F-actin (phalloidin, cyan in merge) around the dendritic cell nucleus in the absence or presence of LPS (overnight stimulation). Orange: DAPI. Yellow arrow indicates the nuclear F-actin cap. G) Quantification of average F-actin intensity above dendritic cell nuclei (at least 10 measurements per condition over 3 donors). H) Airyscan images of immunolabeled vimentin (magenta in merge) around the dendritic cell nucleus in the absence or presence of LPS. I) Quantification of average vimentin intensity above dendritic cell nuclei (at least 10 measurements per condition over 3 donors). J) Example images of dendritic cells cultured on micropatterned surfaces. K) Quantification of the Z-projected nuclear areas of dendritic cells cultured on areas of ≥50 microns, or ≤40 microns shows that a larger adhesion area drives an increase in X-Y nuclear size (minimum of 52 measurements over 4 biological repeats). L) Confocal micrographs of dendritic cells cultured briefly (1 hour) with either cytochalasin D (Cyt; 3 different concentrations) or latrunculin B (LtB). M) Quantification of Z-projected dendritic cell nuclear areas with different actin polymerization inhibitors (minimum of 61 measurements per condition over 4 biological repeats). Data information: Scale bars indicate 20 microns. Statistical significance calculated using a 2-sided unpaired t-test for 2 condition experiments or ANOVA/Tukey multiple comparison test for 3+ condition experiments (with test selected according to distribution pattern of the data) *, P < 0.05; ***, P < 0.001; n.s., not significant. Error Bars= SEM.

In order to confirm the contribution of actin-dependent adhesion on the shape of the nucleus, we cultured dendritic cells on micropatterned substrates with defined adhesion areas. We were able to observe a clear distinction in *Z*-projected nucleus size between cells cultured on adhesion areas of ≥50μm compared to those forced to adhere on areas of ≤40μm (**Fig. 2J and 2K**). We further confirmed these results through culturing dendritic cells in the presence of the actin polymerization inhibitors latrunculin B or cytochalasin D for one hour (**Fig. 2L and 2M**). Both these inhibitors reduced the *Z-*projected nuclear area of dendritic cells to a similar extent.

We also verified the effect of dendritic cell activation on the expression and/or localization of lamin-A/C, which has been shown to alter with tissue stiffness (Swift *et al*., 2013) (Nava *et al*., 2020), and has been shown to turnover less rapidly in activated dendritic cells (Turan *et al*., 2019). Using confocal microscopy we were unable to observe a difference in lamin-A/C distribution in response to LPS stimulation (**Fig. S3E and S3F**). Through western blotting we observed that Lamin-C is the predominant isoform expressed in monocyte-derived dendritic cells, which showed a slight, non-significant reduction in expression in response to LPS stimulation (**Fig. S3G and S3H**). Therefore, we also examined the intensity of Lamin-A/C staining at the nucleus using confocal microscopy (using a set laser power and exposure) (**Fig. S3I**). Through this we observed that lamin-A/C levels reduce in the nucleus (which may be too subtle to observe via western blot), in response to LPS-mediated activation. Therefore, this reduction in lamin-A/C levels may further contribute to the reduction in force observed across Nesprin-2G-based LINC complexes. Indeed, consistent with this we observed that overexpression of Lamin-A/C in dendritic cells is sufficient to block the reduction in *X-Y* nuclear size that accompanies dendritic cell activation ((**Fig. S3J and S3K**).

Given that activation reduces dendritic cell adhesion which drives nuclear spherical deformation, we examined the biophysical properties of dendritic cell nuclei to test if these changes can account for enhanced migration through 2-micron pores. Mechanical stimulation of cells is known to alter the biophysical properties of associated membranes(Hetmanski *et al*., 2019)(Nava *et al*., 2020). However, it is unclear how this would change in dendritic cells, as although activated dendritic cells are less adhesive, the activation process is extremely complex and may involve additional modifications to the nucleus. We examined the fluidity of the dendritic cell membrane using the molecular rotor BODIPY-C10 FRET-FLIM dye (**Fig. 3A and 3B**) (Mika *et al*., 2016), analyzing the lifetime of the dye at the nuclear rim, in a manner akin to described for skin epidermis progenitor cell (EPC) nuclear membranes (Nava *et al*., 2020). This suggested that the fluidity of the nucleus increases in response to LPS stimulation, and thus may promote the flow of the nuclear envelope through gaps in the ECM.

**Figure 3.**
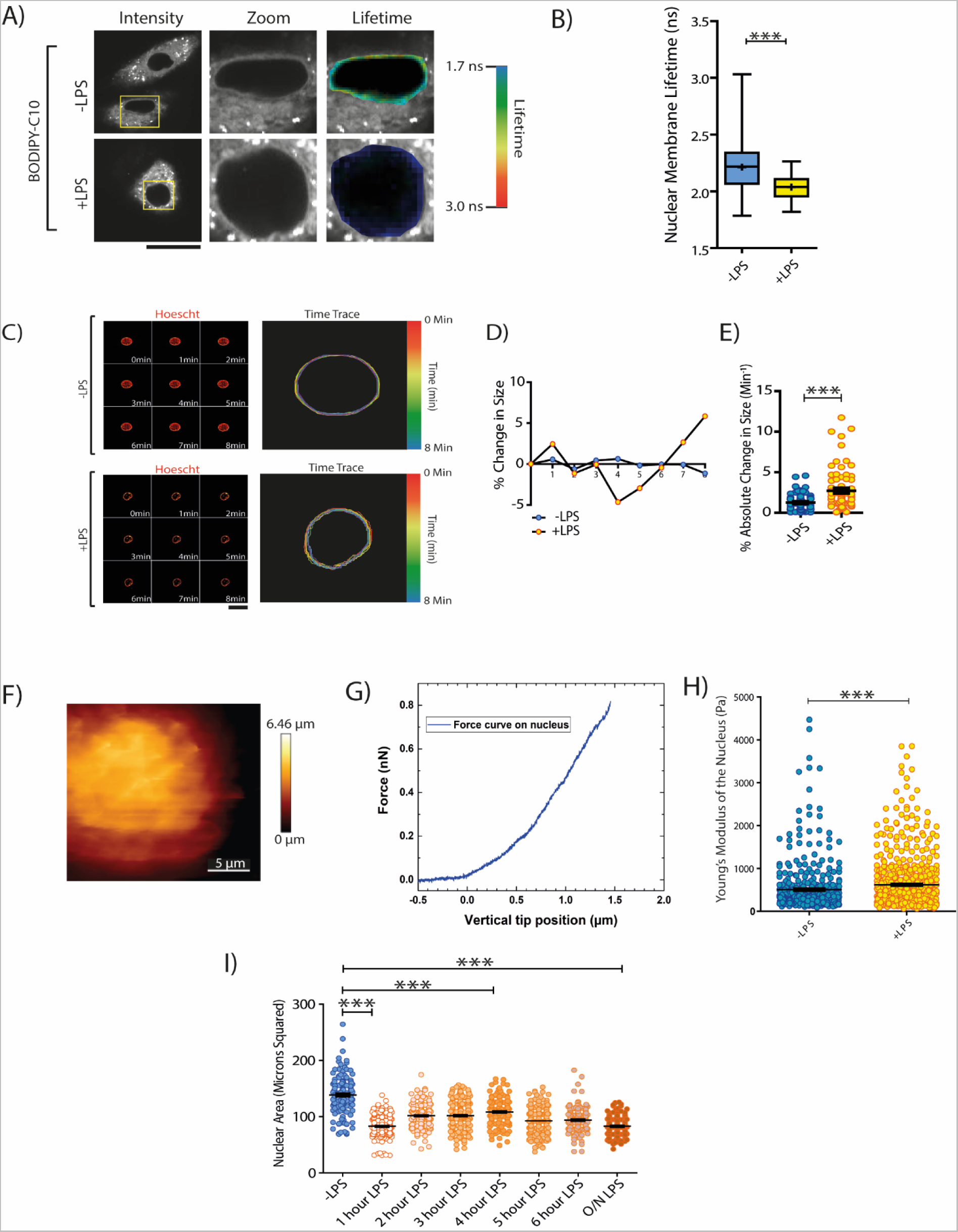
LPS-Stimulation Increases the Stiffness of the Bulk Nucleus. A) Airyscan movies of nucleus in dendritic cells in the absence or presence of LPS (overnight stimulation). B) Example traces of nuclear size fluctuations for dendritic cells in absence (blue) or presence (yellow) of LPS. C) Quantification of absolute percentage changes in nuclear size over time, demonstrating that LPS stimulation increases the temporal size fluctuations of the dendritic cell nucleus (minimum of 67 measurements per condition over 3 biological repeats). D) Intensity and fluorescence lifetime examples of dendritic cells stained with BODIPY-C10, in the presence or absence of LPS. E) Quantification of BODIPY-C10 fluorescence lifetime at the nuclear envelope (minimum of 35 measurements over 4 biological repeats). F) Scanning AFM image of dendritic cell nuclear pouch (a region in which the nucleus is stored). G) Example AFM indentation trace on dendritic cell nucleus. H) Young’s modulus of dendritic cell nuclei quantification across all 4 donors used in AFM experiments (at least 5 cells per donor). Average value for –LPS= 506 Pa, average value for +LPS= 621 Pa. Error bars indicate standard error of the mean (-LPS= 28 Pa, +LPS= 24 Pa). I) Time-course data showing how the dendritic cell Z-projected area changes over time after LPS, starting at 1 hour (at least 145 measurements per condition over 3 biological repeats). Data information: Scale bars indicate 20 microns unless otherwise stated. Statistical significance calculated using 2-sided unpaired t-tests for 2 condition experiments or ANOVA/Tukey multiple comparison tests for 3+ condition experiments (with test selected according to distribution pattern of the data). A Kolmogorov-Smirnov test was used for AFM data to account for the large number of measurements. **, P < 0.01; ***, P < 0.001. Error Bars= SEM. For Box and Whisker plots, Box represents 25th to 75th percentiles, whiskers represents maximum and minumum values, middle band represents data median,+ represents data mean.

To examine bulk nuclear dynamics, we live imaged the dendritic cell nucleus, using airyscan microscopy with the DNA stain Hoechst, in the absence or presence of LPS (**Fig. 3C – 3E, movies S1 and S2**). This revealed that the genome within the nucleus fluctuates in physical distribution more rapidly in activated dendritic cells, compared to inactive dendritic cells. This dynamic behavior may explain how the activated dendritic cell is better able to pass the bulk nuclear content through small gaps in the ECM. However, based on this data it is unclear if the nucleus is becoming more deformable (i.e. softer) or if it is being actively remodeled by the actin cytoskeleton.

In order to test if the dendritic cell nucleus becomes more deformable in response to inflammatory activation we performed atomic force microscopy on dendritic cells to biophysically characterize the deformability of the bulk nucleus (i.e., the nuclear membrane and the chromatin) (Piontek and Roos, 2018). This methodology has previously been used to characterize the dendritic cell plasma membrane (Blumenthal *et al*., 2020) as well as the nuclei of numerous cell types, including epithelial cells and fibroblasts (Rigato *et al*., 2015; Woroniuk *et al*., 2018; Nava *et al*., 2020). Counterintuitively our AFM measurements showed that stimulating dendritic cells with LPS drives bulk nuclear stiffening (**Fig. 3F – 3H**). Therefore, based on these data we conclude that whilst activation induced de-adhesion is sufficient to alter the nucleus shape, and may enhance migration through small gaps in the extracellular matrix (e.g. through altering the biophysical properties of the nuclear membrane), additional force must be required to better squeeze the entire nucleus through gaps in the extracellular matrix, compared to inactive dendritic cells. Indeed this is consistent with previous work that has shown that migration of dendritic cells through small gaps in the ECM depends on myosin activity to squeeze the nucleus (Lämmermann *et al*., 2008; Barbier *et al*., 2019). However it is not clear how this myosin activity is controlled by inflammatory activation as a reduction in cellular adhesion is typically associated with reduced contractility. Therefore we performed phosphoproteomics to elucidate novel changes to the actin cytoskeleton that may facilitate this squeezing.

In order to select relevant time-points of interest for (phospho)proteomic analysis, we first performed a time-course experiment. To do this we examined the shape of dendritic cell nuclei every hour for 6 hours following LPS stimulation (and following overnight stimulation). The time course revealed that the nucleus can shrink in the projected *X-Y* plane within one hour of LPS stimulation (i.e., the shortest time interval addressed), suggesting that the nucleus changes morphology in a (at least partially) transcription-independent manner (**Fig. 3I**). Surprisingly, the size/shape of the nucleus continued to change over 6 hours, partially restoring its projected size 4 hours post-stimulation, before full deformation at the 6-hour mark. Therefore, 1 and 4 hour post-stimulation are ideal time points for identifying key signaling events that drive nuclear deformation; both time-points are deformed relative to unstimulated dendritic cells, yet have slightly different shapes. Thus by selecting these time points we could identify stable signaling events that drive changes to the dendritic cell nucleus at both 1 hour and 4 hours, relative to inactive dendritic cells.

To identify novel early signaling events responsible for the observed changes of the actin cytoskeleton following LPS stimulation, we performed time-resolved MS-based proteomics and phosphoproteomics on dendritic cells (**Fig. S4A, Table S2 and S3**). We compared the proteomes and phosphoproteomes of unstimulated dendritic cells to those stimulated for 1 and 4 hours with LPS. The experiment was performed in six biological replicates (six different donors). Principal component analysis showed that samples clustered per condition, both analyzing the proteome and the phosphoproteome, indicating the reproducibility of the experiment (**Fig. S4B**). On the phosphoproteome level, component-1 explained the difference between unstimulated and LPS-stimulated DCs, while on component-2 the differences between the time points are observed. On the full proteome level, the 1h-treatment condition resembled the unstimulated cells, while the 4h was the most different, consistently with phosphorylation being the fastest cellular event in the TLR4 signaling cascade. Proteomics revealed that 312 proteins, out of 5,593 quantified, significantly changed in expression level over the time-course analyzed (one-way ANOVA, FDR < 0.01) (**Fig. S4C**). These proteins were involved in interferon signaling, cell adhesion, motility and migration (**Fig. S4D**). Differential expression analysis revealed 2,068 significantly regulated phosphosites (one-way ANOVA, FDR < 0.01). To systematically prioritize phosphosites for biological validation, we filtered out sites that were unlikely to have high biological relevance by filtering based on their “functional score”(Franciosa, Martinez-Val and Olsen, 2020). This led to a list of 892 potential phosphosite candidates, which were used to perform hierarchical clustering to determine the directionality of the regulation upon LPS stimulation (**Fig. 4A**). This analysis identified seven expression clusters. Gene ontology enrichment and pathway analysis on each expression cluster revealed that the major biological functions associated with LPS stimulation were related to inflammation (e.g., Toll-like receptor signaling pathway and activation of immune response) and cytoskeleton (e.g., actin filament and focal adhesion), enhancing our confidence in the dataset (**Fig. 4B**). As we observed nuclear deformation at both 1 and 4 hours post LPS stimulation, we specifically focused on cluster F, which contained “stable responders” phosphosites, up-regulated by LPS both at the 1 and 4 hour time points. One of the most significantly enriched terms in this cluster was the gene ontology “Cell junction”. To identify proteins potentially responsible for the observed phenotype, we performed a functional protein network of the cluster F phosphoproteins belonging to this GO term by using STRING (**Fig. 4C; Table S1**) (Szklarczyk *et al*., 2019). This analysis highlighted the most connected nodes (highest degree and network centrality). We focused on cofilin-1 (encoded by the *CFL1* gene - previously shown to regulate nuclear morphology (Baarlink *et al*., 2017)), which was phosphorylated on the serine 41 upon LPS (**Fig. 4D to 4F, Table S1**). However it is likely that also other members of this cluster regulate changes to dendritic cell adhesion/actin dynamics upon inflammatory activation.

**Figure 4.**
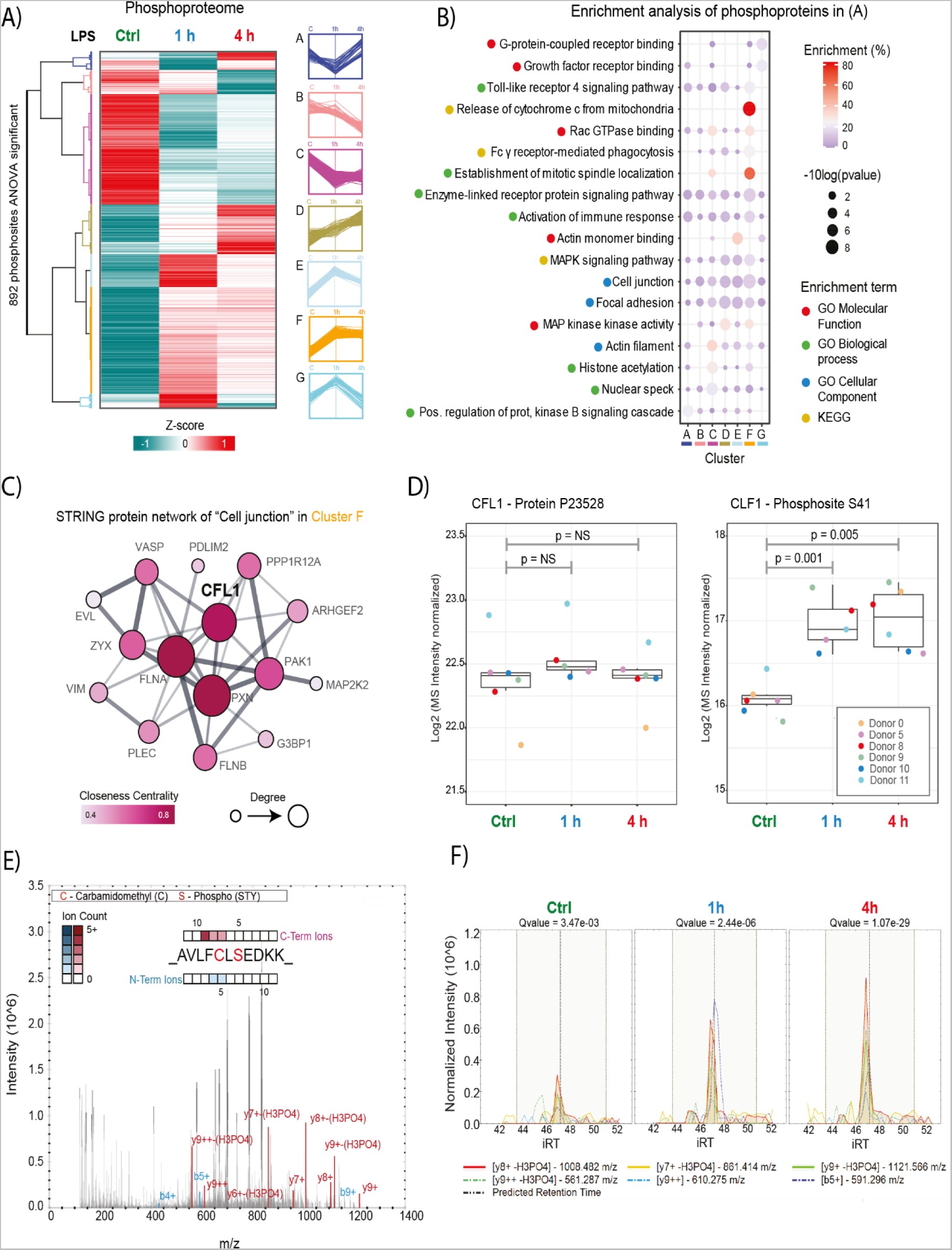
Phosphoproteomic Analysis of Dendritic Cell Inflammatory Activation. A) Hierarchical clustering analysis by using Pearson correlation distance of phosphosite intensities. Values were normalized, imputed, batch-corrected and scaled (Z-score) before clustering. Only sites with one-way ANOVA FDR < 0.01 and functional score > 0.45 were used for clustering. N = 6 donors. B) Gene ontology and Pathway (KEGG and Reactome) enrichment analysis on each cluster shown in A. Enrichment was performed on phosphoprotein level against the entire proteome quantified. C) Functional STRING protein network of the phosphoproteins belonging to Cluster F and to the GO “Cell junction”. D) Boxplot of the normalized MS signal, after Log2 transformation, for the protein cofilin-1 (left) and the phosphorylated serine 41 of cofilin-1 (right). P values were calculated by two-sided paired Student’s t-test. E) MS/MS spectra for the cofilin-1 phosphopeptide containing the phosphorylated serine 41. F) Extracted MS2 ion chromatogram for the cofilin-1 phosphopeptide shown in E. Each line represents a fragment ion. iRT = independent retention time (in minutes). Data information. For Box and Whisker plots, Box represents 25th to 75th percentiles, whiskers (when displayed) represents maximum and minumum values, middle band represents data median.

To determine whether cofilin-1^pS41^ drives nuclear deformation, we overexpressed GFP-tagged wildtype (WT) cofilin-1, along with phosphomimetic (S41E) and phosphodead (S41A) cofilin-1 mutants in dendritic cells. Analysis of the projected *X-Y* size of nuclei revealed that phosphomimetic cofilin-1 (S41E) can induce deformation of the dendritic cell nuclei (**Fig. 5A and 5B**), although it is unlikely that cofilin-1 uniquely regulates deformation of the dendritic cell nucleus. We also verified that cofilin-1 S41E and S41A were not altering the shape of the nucleus through reducing dendritic cell spreading. Indeed, although the nuclei reduced in *X-Y* projected size, the *X-Y* projected size of the entire cell did not reduce (**Fig. S5A**). In fact overexpression of S41E cofilin-1 lead to a small (non-significant) increase in cell spreading, relative to WT cofilin-1. Therefore, is seems likely that cofilin-1^pS41^ contributes to nuclear deformation but not to the de-spreading process. Thus, although nuclear deformation is driven by reduced dendritic cell adhesion, it would seem that changes in cell spreading do not entirely account for dendritic cell nuclear deformation. Surprisingly, the phosphodead mutant was also able to induce deformation of the dendritic cell nucleus in the *X-Y* plane. The reasons for this are unclear. However, molecular simulations with enhanced sampling techniques indicated that the structure of cofilin-1 is very stable; phosphorylation of serine 41 (and dephosphorylation of serine 3) leaves the structure relatively unchanged (**Fig. S5B – S5D**). Therefore, our data suggest that phosphorylation of cofilin-1 at serine 41 may instead alter cofilin-1’s interactome, through loss of the OH group on the serine (as opposed to altering cofilin-1’s molecular structure). However confirming this will require further experimentation. We conclude that cofilin-1^pS41^ promotes dendritic cell nuclear deformation, but (based on our phosphoproteomic data) is unlikely to be the sole contributor to this phenotype.

**Figure 5.**
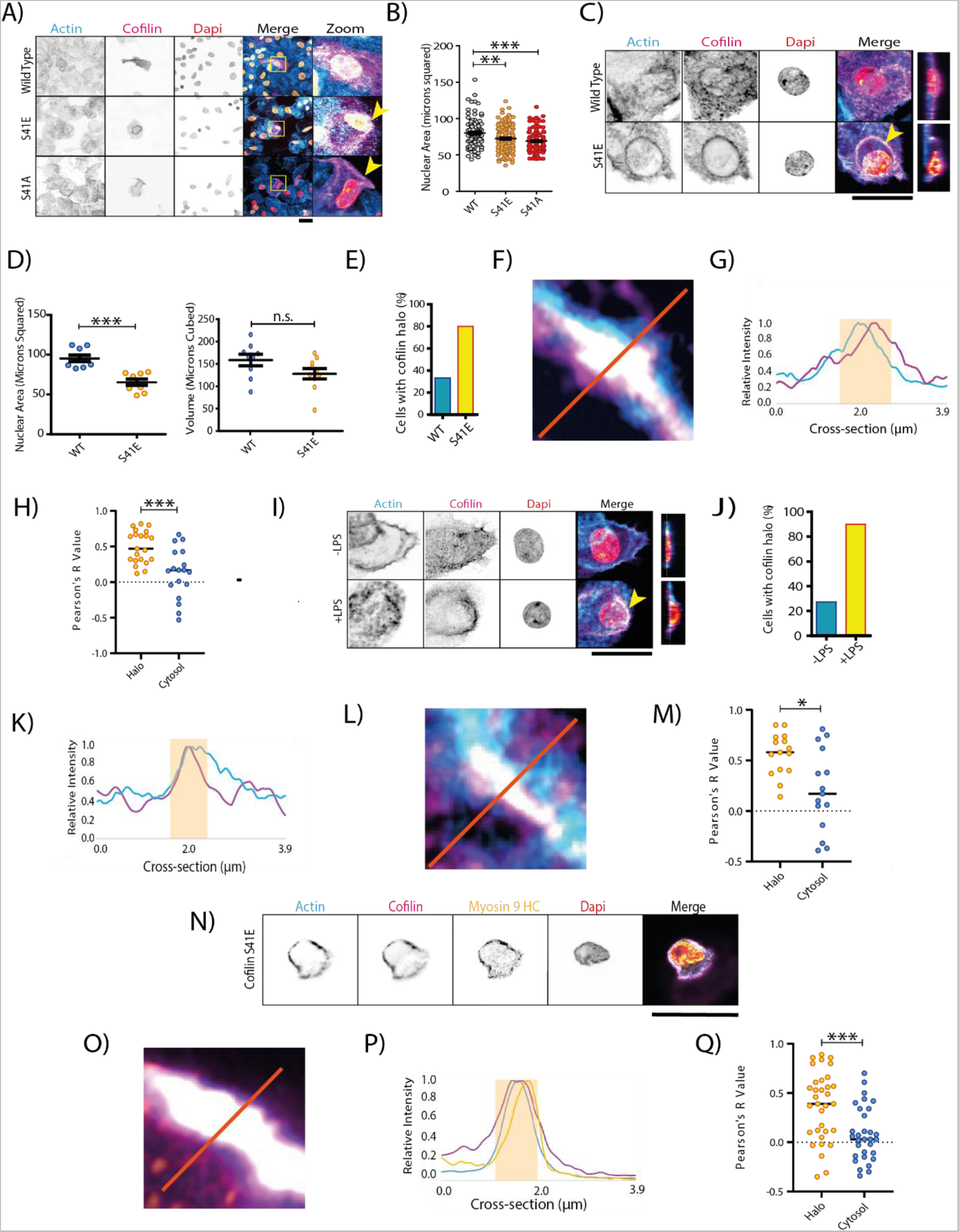
Nuclear Deformation is driven by Cofilin-1 S41 Phosphorylation. A) Confocal micrographs of dendritic cells expressing either WT, S41E or S41A cofilin-1 fused to GFP (magenta in merge). Cyan: phalloidin. Orange: DAPI. Yellow arrows indicate qualitatively observed peri-nuclear cofilin-1 halos. B) Quantification of the Z-projected nuclear area of dendritic cells expressing cofilin-1 variants (data for at least 108 measurements per condition over 3 biological repeats). C) Airyscan images of dendritic cells expressing WT or S41E cofilin-1 variants. Yellow arrow indicates peri-nuclear cofilin-1 halo. D) X-Y projected area along with volume of the same nuclei of cells expressing either WT cofilin-1 or S41E cofilin-1. E) Quantification of the number of cells with cofilin-1 halos around the nucleus (expressing WT or S41E cofilin-1 observed in airyscan images (minimum of 9 cells per condition over 3 biological repeats). F) Example of cofilin-1 - F-actin co-localization in a dendritic cell expressing cofilin-1 S41E. G) Line profile of normalized cofilin-1 (purple) and F-actin (blue) intensity as indicated in panel F. Highlighted region indicates area of substantial overlap. H) Pearson’s correlation coefficient of cofilin-1 – F-actin at the halos and in the cytosol of cells expressing cofilin-1 S41E (minimum of 18 measurements over 3 biological repeats). 3 regions per compartment per cell were analyzed. I) Airyscan images of cofilin-1 and F-actin in dendritic cells cultured overnight in the presence or absence of LPS (overnight stimulation). Yellow arrow indicates peri-nuclear cofilin-1 halo. J) Quantification of the percentage of cells with cofilin-1 halos around the nucleus (cultured in the presence or absence of LPS), observed in airyscan images (minimum of 10 cells per condition over 3 biological repeats). K) Example of cofilin-1 – F-actin co-localization in dendritic cells cultured overnight with LPS. L) Line profile of normalized cofilin-1 (purple) and F-actin (blue) intensity as indicated in panel K. Highlighted region indicates area of substantial overlap. M) Pearson’s correlation coefficient of cofilin-1 – F-actin at the halos and in the cytosol of cells cultured overnight with LPS that displayed F-actin-cofilin-1 bundling (minimum of 15 measurements over 3 biological repeats). N) Airyscan images of dendritic cells S41E cofilin-1 stained for F-actin and Myosin 9 HC. O) Example of S41E cofilin-1 - F-actin-myosin co-localization in a dendritic cell expressing cofilin-1 S41E. P) Line profile of normalized cofilin-1 (purple), F-actin (blue) and myosin 9 HC (yellow) intensity as indicated in panel P. Highlighted region indicates area of substantial overlap. Q) Pearson’s correlation coefficient of cofilin-1 – myosin 9 HC at the halos and in the cytosol of cells expressing cofilin-1 S41E (minimum of 31 measurements over 3 biological repeats Data information: Scale bars indicate 20 microns. Statistical significance calculated using 2-sides unpaired t-tests for 2 condition experiments or ANOVA/Tukey multiple comparison test for 3+ condition experiments (with test selected according to distribution pattern of the data). A 2-sided paired t-test was used for panels H and M. **, P < 0.01; ***, P < 0.001. Error Bars= SEM.

We also examined the impact cofilin-1 S3 phosphorylation on *Z-*projected nuclear size. Based on previous findings (Verdijk *et al*., 2004), cofilin-1^pS3^ is dephosphorylated upon LPS stimulation, which has been reported to regulate nuclear morphology (Baarlink *et al*., 2017). De-phosphorylation at this residue enables its actin severing activity (Bravo-Cordero *et al*., 2013). However, at both 1 and 4 hours, this was a non-significant change in our phosphoproteomic data (**Fig. S4E**). Furthermore, overexpression of a cofilin-1 S3A mutant (phosphodead) was not sufficient to induce the nuclear membrane deformation in dendritic cells (**Fig. S5E and S5F**) (Garvalov *et al*., 2007). Phosphorylation of cofilin-1 at serine 41 has been previously observed in several phospho-proteomic studies, including in mouse dendritic cells and human cancer cells (Dephoure *et al*., 2008; Kettenbach *et al*., 2011; Klammer *et al*., 2012; Zhou *et al*., 2013; Shiromizu *et al*., 2013; Sharma *et al*., 2014; Dally *et al*., 2006; Mertins *et al*., 2013; Mertins *et al*., 2014; Mertins *et al*., 2016; Mertins *et al*., 2017). However, the functional relevance of this phosphorylation has not been previously studied.

In order to further understand the contribution of cofilin^pS41^ to the deformation of the cell nucleus, we analyzed dendritic cells overexpressing WT or S41E cofilin-1, using Airyscan microscopy. First, we verified that the nuclei were deforming and not shrinking (**Fig. 5C and 5D**), as we observed for activated dendritic cells (**Fig. 1C and 1D**). This revealed that although S41E cofilin-1 can significantly reduce the *X-Y* projected size, it does not significantly change the nuclear volume; the nuclei extend into the Z-plane similar to our observations with LPS activation. Furthermore, qualitative observation of cofilin-1 S41E (as well as qualitative observation of S41A expressing dendritic cells) suggests that it (and therefore cofilin-1^pS41^) accumulates around the nucleus (**Fig. 5A**). Therefore, we examined the nature of these structures using Airyscan microscopy. This revealed that the majority of cells overexpressing cofilin-1 S41E, with deformed nuclei form a cofilin-1 halo (**Fig. 5C and 5E**). The majority of these halos co-localized with F-actin (**Fig. 5F – 5H**). We also noticed that F-actin filaments are lost from the nuclei in dendritic cells expressing both WT and S41E cofilin. These are likely lost as an overexpression artifact. Using the mN2G-TS FRET probe, we confirmed that there is no difference in force across actin-based LINC complexes in cells expressing WT or S41E cofilin (**Fig. S5G and S5H**). Therefore, S41E cofilin is able to remodel the nuclei through a mechanism that does not involve actin-based LINC complex remodeling. It seems possible therefore, that the observed cofilin halos are not passively sequestering actin away from the nucleus, but may instead be driving an active remodeling process. Indeed, Airyscan microscopy of endogenous cofilin-1 confirmed that a similar halo forms in the majority of LPS stimulated cells with highly deformed nuclei (stimulated overnight at cofilin S41 phophorylation is a stable responder in the phosphoproteomic dataset) (**Fig. 5I and 5J**). We were able to observe co-localization between cofilin-1 and F-actin in approximately 50% of activated cells (**Fig. 5K – 5M**). Cofilin-1-based actin clustering has been previously observed with purified proteins (Pfannstiel *et al*., 2001) in *Drosophila* (Wu *et al*., 2016), rats (Hylton *et al*., 2021) and pathologically, in Alzheimer’s disease (Minamide *et al*., 2000), however this is the first example of a direct cofilin-1 phosphorylation that can drive cofilin-1-based actin clustering, although the precise role of these halos will require further investigation. We therefore hypothesized that these halos may be contractile, therefore we stained for myosin 9 heavy chain in cells expressing mCherry-cofilin S41E. This revealed a strong co-localization between cofilin S41E and myosin motors (**Fig. 5N-5Q**). We have therefore termed these halos Cofilin-ActoMyosin (CAM)-rings.

In order to test the impact of cofilin S41 phosphorylation on nuclear dynamics and migration we first analysed the impact of overexpressing WT or S41E cofilin on nuclear shape fluctuations (**Fig. 6A-6D; movies S5 and S6**). This revealed that the shape of the nucleus fluctuates more in cells expressing phosphomimetic cofilin (as with LPS activated dendritic cells vs inactive dendritic cells). Then, to test the impact of cofilin phosphorylation on migration in 3D environments, we overexpressed GFP-WT and RFP-S41E cofilin in activated dendritic cells, mixed the populations, and seeded them onto bovine dermal based collagen gels (produced at 1.7 mg.ml^−1^). (**Fig. 6E – 6I; movies S7 and S8**). This revealed that overexpression of cofilin-S41E is sufficient to increase the MSD in active dendritic cells (**Fig. 6J and 6K**). These results demonstrate that phosphorylation of cofilin at serine 41 is sufficient to mimic the increase in morphological dynamism associated with dendritic cell activation and simultaneously reduces the relative confinement exerted on dendritic cells by the extracellular matrix. As these collagen gels have holes approaching the physical limit of migration (as determined by the nucleus (Wolf *et al*., 2013)), we conclude that cofilin^pS41^ driven nuclear remodeling facilitates efficient dendritic cell migration through the ECM, likely through CAM-ring mediated contractility.

**Figure 6.**
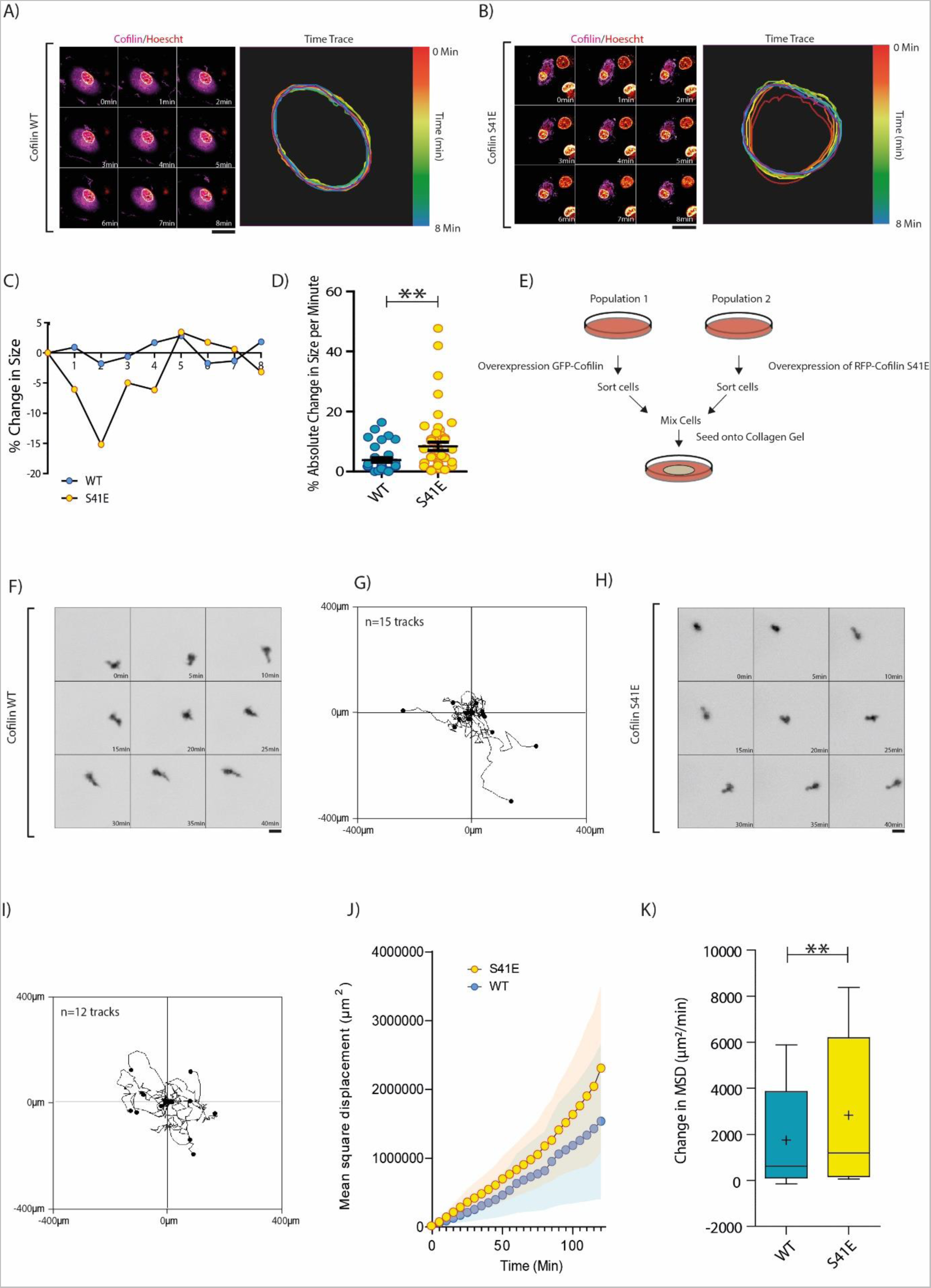
S41 Phosphorylation of Cofilin Increases Migration in Confined Environments. A) Airyscan movie and time trace of nucleus in dendritic cell overexpressing WT cofilin. B) Airyscan movie and time trace of nucleus in dendritic cell overexpressing cofilin S41E. C) Example traces of nuclear size fluctuations for dendritic cells overexpressing WT cofilin (blue) or cofilin S41E (yellow). D) Quantification of absolute percentage changes in nuclear size over time, demonstrating that phosphorylation of cofilin at serine 41 increases the temporal size fluctuations of the dendritic cell nucleus (minimum of 67 measurements over 3 biological repeats). E) Schematic demonstrating migration experiment setup. F) Example movie of activated dendritic cell overexpressing WT cofilin. G) Examples migration traces of activated dendritic cell overexpressing WT cofilin. H) Example movie of activated dendritic cell overexpressing S41E cofilin. I) Examples migration traces of activated dendritic cell overexpressing S41E cofilin. J) Example MSDs of activated dendritic cells overexpressing WT or S41E cofilin. Shaded region represents SEM. K) Quantification of rate of MSD change per donor, for activated dendritic cells overexpressing WT or S41E cofilin (75 measurements per condition over 3 biological repeats) Data information: Statistical significance calculated using 2-sides unpaired t-tests for 2 condition experiments or ANOVA/Tukey multiple comparison test for 3+ condition experiments (with test selected according to distribution pattern of the data). Scale bars indicate 20 microns. **, P < 0.01; ***, P < 0.001. Error Bars= SEM. For Box and Whisker plots, Box represents 25th to 75th percentiles, whiskers represents maximum and minumum values, middle band represents data median,+ represents data mean.

## Discussion

Dendritic cells are found in virtually all tissues, yet they need to migrate to lymph nodes to initiate a novel adaptive immune response (Patente *et al*., 2019). Therefore, dendritic cells must have extremely plastic biochemical and mechanosensitive signaling machinery that can adapt to residency within each tissue, yet override mechanical signals once activated. Indeed, activated dendritic cells are less adhesive (Van *et al*., 2006; West *et al*., 2008) and integrin-free dendritic cell migration is both viable and efficient (Lämmermann *et al*., 2008b; Reversat *et al*., 2020). However, a perceived disadvantage of adhesion-free migration is that the nucleus blocks migration through small pores (<3-microns), that could otherwise be expanded through ECM remodeling (Wolf *et al*., 2013). Our study shows that dendritic cell activation can (partially) compensate for this, as the nucleus becomes more spherically deformed and is better able to pass through 2-micron gaps in the ECM. These changes to the shape of the nucleus seem to also be partially passive, via a reduction in adhesion, and partially active, with cofilin^pS41^ triggering the assembly of perinuclear CAM-rings. Indeed, the assembly of a contractile ring around the nucleus, in response to LPS stimulation, makes sense of the fact that the dendritic nucleus seems to be able to better pass through gaps in the ECM whilst simultaneously becoming stiffer. However it is not clear how or why dendritic cells would stiffen their nucleus in response to activation. A softer nucleus should flow better through gaps in the ECM and the loss of the lamin-A/C typically drives nuclear softening. We speculate therefore that nuclear stiffening may be driven by epigenetic regulation of the genome. Such regulation has previously been shown to directly control nuclear stiffness (Nava *et al*., 2020)(Wang *et al*., 2018)(Chalut *et al*., 2012) (Stephens *et al*., 2018) and may be essential for the transcriptional changes that underpin dendritic cell activation.

Dendritic cell activation is extremely well understood at the genetic level, with numerous signaling receptors and transcription factors identified (Dalod et al., 2014). However, the complex (de)phosphorylation events underpinning this activation are poorly understood. Although a few high-quality phosphoproteomic studies describing these (de)phosphorylation events in dendritic cells have been produced (Mertins *et al*., 2017; Korkmaz *et al*., 2018; Li *et al*., 2021), only 1 has been published for human dendritic cells(Li *et al*., 2021). Our phosphoproteomic data both contributes to this understanding and provides (to date) the most comprehensive phosphoproteomic description of human myeloid cell activation, having identified 2,068 statically significant (de)phosphorylation events. Indeed, our phosphoproteomic description of dendritic cell activation has revealed a previously unidentified adhesion-associated signaling cluster, focused around cofilin-1^pS41^. Therefore studies of these (de)phosphorylation events will be particularly critical, as the inflammatory response of dendritic cells underpins not only changes to adhesion machinery, but also to the release of inflammatory cytokines and antigen presentation to T cells.

How does cofilin-1^pS41^ drive migration through the ECM? Our data suggests that this is ultimately achieved through force transmission to the nucleus, via the assembly of perinuclear CAM-rings. Indeed, previous work has shown that dendritic cells path-find through the ECM by actively deforming their nucleus to protrude into multiple pores in their immediate vicinity (Renkawitz *et al*., 2019). This allows the dendritic cell to select the largest pore to migrate through. However, it seems likely that the myosin motors on the CAM-rings also drive bulk squeezing of the cytosol, driving efficient migration through the ECM. Furthermore, cofilin-1^pS41^ may, in parallel, perform additional functions that reduce the confining effect of 3D-matrix fibers on dendritic cell migration.

In conclusion, we have demonstrated that dendritic cell activation triggers nuclear deformation and drives enhanced transmigration efficiency through 2-micron pores. Although nuclear deformation partially results from reduced dendritic cell adhesion, we have shown that increased trans-migration efficiency likely requires active, actin-driven deformation of the nucleus, as the bulk nucleus seems to stiffen in response to LPS. Active deformation of the nucleus is achieved by the phosphorylation of cofilin-1^pS41^. This phosphorylation leads to the assembly of nuclear proximal CAM-rings (**Fig. 7A and 7B**), active remodeling of the nucleus and enhanced migration through 3D-ECM. Given that cofilin-1^pS41^ has been observed in numerous additional phosphoproteomic datasets, it seems likely that cofilin may perform similar functions in other cell-migration contexts, such as during wound healing or cancer metastasis.

**Figure 7.**
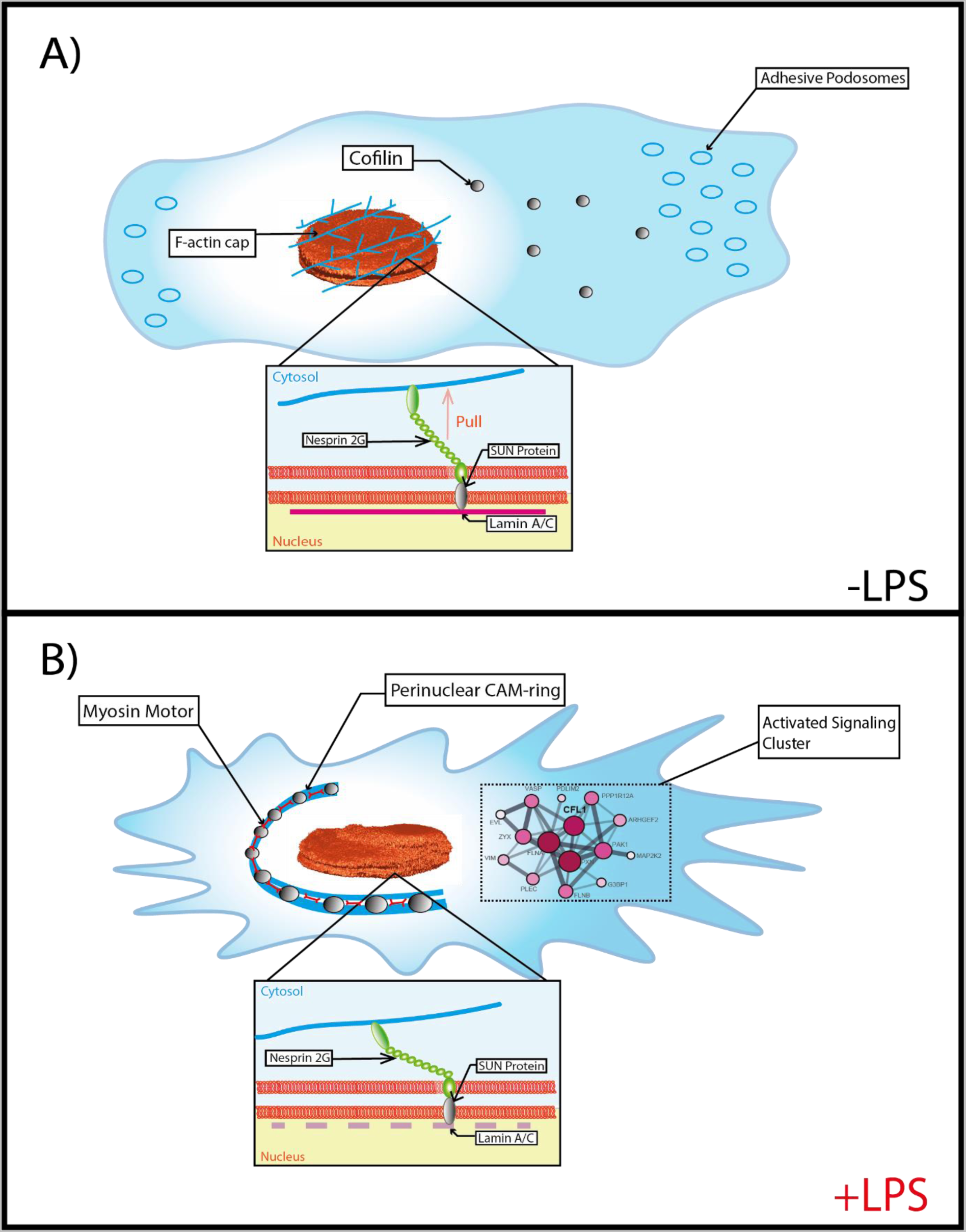
Working Model. A) Model in absence of LPS. Inactive dendritic cells are highly adhered and have a perinuclear actin cap that exerts force on nesprin-2 based LINC complexes and maintains a flat, disc-shaped nucleus in stiffer tissues. B) Part two of the model. When activated through LPS stimulation, cell spreading is reduced resulting is a spherically deformed, raison shaped nucleus, with reduced force across nesprin-2 based LINC complex. Cofilin phosphorylation at serine 41 promotes the formation of CAM-rings (consisting of cofilin, actin and myosin) that exert force (directly or indirectly) on the nucleus, driving it’s passage through gaps in the ECM.

## Materials and Methods

### Cells

Monocyte-derived dendritic cells were obtained by differentiating monocytes with interleukin (IL)-4 (300 µg/ml) and granulocyte-macrophage colony-stimulating factor (GM-CSF; 450 µg/ml) for 6 days in RPMI supplemented with 10% serum, antibiotics (100 μg ml^−1^ penicillin, 100 µg ml^−1^ streptomycin and 0.25 µg ml^−1^ amphotericin B, Gibco), and 2 mM glutamine. Monocytes were isolated from the blood of healthy donors (informed consent and consent to publish obtained, approved by the ethical committee of Dutch blood bank Sanquin) as previously described (de Vries *et al*., 2002). LPS (O111:B4, Sigma Aldrich 32160405) stimulation was carried out overnight unless otherwise stated.

CD14^+^ macrophages were obtained as previously described. In brief CD14^+^ monocytes were isolated using MACs kits (Miltenyi) from the blood of healthy donors. The CD14^+^ monocytes were differentiated for 7 days on low-adherence plates (Corning) in RPMI supplemented with 10% serum, antibiotics (100 μg ml^−1^ penicillin, 100 µg ml^−1^ streptomycin and 0.25 µg ml^−1^ amphotericin B, Gibco), and M-CSF (100ng/ml) for 7 days.

### Reagents Antibodies

The following primary antibodies were used in this study: cofilin-1 (Thermofisher GT567, 1:200): 1:200, lamin-A/C (Abcam ab108595, 1:200 for immunofluorescence, 1/1000 for western blot), vimentin (Abcam ab92547, 1:200), tubulin (Novus Biologicals YOL1/34), GAPDH (Cell Signaling 2118, 1:500), Myosin 9 HC (Thermofisher 5D9D2, 1:200).

The following secondary antibodies and reagents were for immunofluorescence: donkey anti-rabbit 647 (Thermo Fisher A31573), goat anti-rat 488 (Thermo Fisher A11006), donkey anti-mouse 647 (Thermo Fisher A31571), phalloidin Alexa-Fluor-488 (Thermo Fisher A12379), phalloidin Alexa-Fluor-647 (Thermo Fisher A22287), DAPI (Sigma Aldrich 32670).

### Total Cell Lysis and Western Blotting

Total cell lysates were obtained using a boiling SDS lysis buffer (5% sodium dodecyl sulfate (SDS), 5 mM tris (2-carboxy-ethyl)phosphine (TCEP), 10 mM chloroacetamide (CAA), 100 mM Tris, pH 8.5). Lysates were then incubated for 10 minutes at 95°C. Lysates were resolved on polyacrylamide gels and transferred to methanol-activated PVDF. Blots were scanned with an Odyssey XF imaging system (LI-COR Biosciences).

### Confocal Microscopy

Cells were seeded onto glass coverslips for at least 12 hours before fixation. Cells were fixed in 4% PFA for 15 minutes at room temperature, or ice-cold methanol for 10 minutes at −20°C (for lamin-A/C immunolabeling). Cells fixed in PFA were permeabilized for 5 minutes in a 0.1% (v/v) Triton-X100 solution. Cells were blocked in a 20 mM glycine 3% BSA PBS-based solution for 1 hour before antibody staining.

Transient transfection of dendritic cells was achieved using the Neon-transfection system (Thermo Scientific). In brief, 1.2 million cells were washed with PBS and suspended in 115 µL of buffer R with 5 µg of DNA. Cells were pulsed twice for 40 ms at 1000 V. Cells were then transferred to phenol red-free RPMI with 20% serum for at least 4 hours before imaging. Live cell imaging was performed in OptiKlear solution (Abcam ab275928).

Images were collected with a Zeiss LSM 800 microscope equipped with a Plan-Apochromat (63x/1.4) oil DIC M27 (FWD=0,19 mm) objective (Zeiss). Images were acquired using the ZEN software (version 2.3). DAPI was excited by a 405 laser, and Alexa Fluor 488 phalloidin was excited by a 488 laser. For Z-series, a slice interval of 0.31 µm was used. Airyscan microscopy was performed with a Zeiss LSM 800 airyscan microscope. A Z-interval of 0.17 µm was used. Images were acquired using the ZEN software (version 2.3). Images were subject to airyscan processing following acquisition. 3D reconstruction was achieved using the IMOD software package (Kremer, Mastronarde and McIntosh, 1996).

### ECM coating of Glass Coverslips

Glass coverslips were coated as previously described. In brief, sterile glass coverslips were coated in a PBS solution containing each respective ECM protein (collagen (0.3 mg ml^−1^), fibronectin (33 µg ml^−1^), laminin (0.1-0.2 mg ml^−1^), vitronectin (0.1 mg ml^−1^), fibrinogen (0.1 mg ml^−1^)) overnight. The coverslips were then washed with PBS before the seeding of dendritic cells in RPMI supplemented with serum and antibiotics.

### Inhibitors

Cytochalasin D (Merck C2618) was used at concentrations ranging from 100nM to 10 µM, Latrunculin B (Abcam Ab144291) was used at 100nM. SMIFH2 (Merck S4826) was used at 10 µM, CK666 (Merck SML0006) was used at 150 µM, Blebbistatin (Merck B0560) was used at 50 µM. All inhibitors were added for 1 hour.

### Bodipy-C10

The BODIPY C10 dye (kind gift from Dr. Ulf Diederichsen, Georg-August-Universität Göttingen, Germany) was used at 4 μM to stain cells for 30 minutes before imaging. Cells were then washed twice with phenol-red free RPMI before imaging. Cells were imaged using a PicoQuant MicroTime 200 microscope equipped with an Olympus (100x/1,4) oil immersion objective. Images were acquired using the SymPhoTime 64 software. Data analysis of FLIM images was performed using the open-source FLIMfit software (version 5.1.1.). For analyzing the nuclear membrane, parts of the respective membrane were selected to avoid adjacent organelles.

### Atomic Force Microscopy

AFM measurements were performed using a commercial JPK NanoWizard mounted on an inverted optical microscope (Olympus). The AFM was equipped with a JPK Biocell heated stage and the measuring temperature was kept at a constant 37°C. Silicon nitride cantilevers with a triangular pyramidal probe and a nominal spring constant of 0.1 N/m were used (MLCT-BIO E, Bruker). The cells were seeded on clean glass coverslips. Cell nuclei were identified through light microscopy and then imaged in Quantitative Imaging mode before indenting. Indentations were performed by drawing a 6×6 points grid in a 3×3 µm area on the highest point of the nucleus and then indenting each point once, after which moving on to a different cell. Indentations were performed to a set-point of 1 nN at a velocity of 1 µm/s. In between cells, a glass curve was taken to check for tip contamination.

Both image and force curves were processed using JPK Data Processing software. Curves were fit using the inbuilt Hertz / Sneddon model for triangular pyramid-shaped probes to obtain the Young’s modulus. In Fig 3H each data point corresponds to a single indentation of the used 6×6 grid. Batch processing was used with manually chosen fitting ranges. Typical deformation of the cells at the place of the nucleus was ~1.5 µm. This indicates that the indentation was much deeper than the actin cortex layer which is several hundreds of nanometer thick (Roos *et al*., 2003; Clausen *et al*., 2017) and that a considerable part of the nucleus was deformed.

### Flow Cytometry

Monocyte-derived dendritic cells and monocytes isolated with the CD14 MACS kit from Miltenyi biotec (130-050-201) were collected in a V-bottom plate at 10^5^ cells/well (Thermo scientific # 10462012) and pelleted (300 *xg*, 3 min, 4°C). Next, cells were blocked in 2% human serum and PBS for 30 min at 4°C and stained with DC-SIGN (Beckman Coulter #A07407), CD80 (BioLegend #305214), CD86 (BD #555658) and HLA-DR (BD #559866) FACS-Abs also in human sera with PBS for 60 min at 4°C. After staining cells were washed in PBS two times, recorded on the CytoFlex S (Beckman Coulter), and analysed with FlowJo.

### Ratiometric FRET Analysis

Ratiometric FRET images were obtained from moDCs expressing mN2G-TS (gift from Daniel Conway; Addgene 68127) or pcdna nesprin HL (gift from Daniel Conway; Addgene 68128) (Arsenovic and Conway, 2018)Ratiometric FRET images were analysed using an in-house ImageJ macro. In brief the YFP-FRET signal was divided by the sum of the YFP-FRET and CFP signal.

### Plasmids

The following plasmids were used in this study: mN2G-TS (gift from Daniel Conway; Addgene 68127) (Arsenovic and Conway, 2018), pcdna nesprin HL (gift from Daniel Conway; Addgene 68128), pEGFP-N1 human cofilin-1 WT (gift from James Bamburg; Addgene 50859), pEGFP-N1 human cofilin-1 S3E (gift from James Bamburg; Addgene 50861), pEGFP-N1 human cofilin-1 S3A (gift from James Bamburg; Addgene 50860) cofilin-1 (Garvalov *et al*., 2007). pEGFP-N1 human cofilin-1 1 S41A and pEGFP-N1 cofilin-1 S41E, pEGFP-C1 Lamin A/C, were generated as synthetic genes (Genscript).

### 3D Cell Culture

Custom-made fibronectin stiffness gels were purchased from 4D Cell. Cells were cultured overnight in corning plastic in the presence or absence of LPS, at 37°C, 5% CO_2_. Cells were then detached via incubation in PBS at 4°C and re-seeded onto stiffness gels, before being allowed to adhere for at least 3 hours before fixation with PFA. Cells were then stained with phalloidin and DAPI, and imaged using confocal microscopy.

### Micropatterned Substrate

Micropatterned slides were purchased from 4D Cell (UM001). Cells were allowed to adhere on the slides prior to being 4% PFA for 15 minutes. Samples were then stained with DAPI and phalloidin, mounted and imaged.

### Gelatin Impregnated Filters

Polycarbonate membrane filters (Sterlitech, Kent, WA: PCT1025100 – 1-micron pores, PCT2013100 – 2-micron pores, PCT3013100 – 3-micron pores) were washed with 70% ethanol before being pressed between a glass coverslip and a parafilm sheet, with a droplet of fluorescent gelatin (DQ Gelatin – Thermo D12054) for 10 minutes. The membranes were then washed with PBS before cells were seeded on top of the membrane filter. Following overnight incubation, samples were fixed in 4% PFA for 15 minutes. Samples were then stained with DAPI and phalloidin, mounted and imaged.

Invasion was calculated by normalizing the total number of cells on the underside of filters to the seeding density on the topside of the filters. This was then normalized for the +LPS condition to the –LPS condition.

### Cell lysis and digestion for proteomics analysis

Cells were lysed in boiling 5% SDS buffer (in 100 mM Tris-HCl pH 8.5, 5 TCEP, 10 mM CAA). Lysates were heated for 10 minutes. After sonication, protein concentration was estimated by BCA assay (Pierce). Protein digestion using the PAC method) (Batth *et al*., 2019)was automated on a KingFisher™ Flex robot (Thermo Fisher Scientific) in 96-well format, as previously described (Leutert *et al*., 2019)(Bekker-Jensen, Martínez-Val, *et al*., 2020). The 96-well comb is stored in plate #1, the sample in plate #2 in a final concentration of 70% acetonitrile and with magnetic amine beads (ReSyn Biosciences) in a protein/bead ratio of 1:2. Washing solutions are in plates #3–5 (95% Acetonitrile) and plates #6–7 (70% Ethanol). Plate #8 contains 300 μl digestion solution of 50 mM ammonium bicarbonate (ABC), LysC in an enzyme/protein ratio of 1:500 (w/w) and trypsin in an enzyme:protein ratio of 1:250. The protein aggregation was carried out in two steps of 1 min mixing at medium mixing speed, followed by a 10 min pause each. The sequential washes were performed in 2.5 min and slow speed, without releasing the beads from the magnet. The digestion was set to 12 h at 37 °C with slow speed. After overnight digestion, enzymatic activity was quenched by acidifying the lysates using trifluoroacetic acid (TFA) at a final concentration of 1% and ensuring the pH of the samples being around 2. Digested peptides for single-shot proteome analysis were loaded directly on C18 evotips (Evosep) for MS analysis. Digested peptides for phosphoproteomics were purified and concentrated on reversed-phase C18 Sep-Pak cartridges (Waters). After elution with 40% acetonitrile (ACN) followed by 60% ACN, a SpeedVac concentrator (ThermoFisher Scientific), operating at 60°C, was utilized to concentrate the samples. Peptide concentration was estimated by measuring absorbance at A280 on a NanoDrop spectrophotometer (ThermoFisher Scientific).

### Enrichment of phosphorylated peptides

Ti-IMAC phosphopeptide enrichment was carried out on a KingFisher™ Flex robot (Thermo Fisher Scientific) in 96-well format, as previously described (Bekker-Jensen, Martínez-Val, *et al*., 2020). 200 μg of peptide were used for enrichments, with 20 μl of magnetic Ti-IMAC HP beads (ReSyn Biosciences). The 96-well comb is stored in plate #1, Ti-IMAC HP beads in 100% ACN in plate #2 and loading buffer (1 M glycolic acid, 80% ACN, 5% TFA) in plate #3. The sample is mixed with loading buffer and added in plate #4. Plates 5–7 are filled with 500 μl of washing solutions (loading buffer, 80% ACN, 5% TFA, and 10% ACN, 0.2% TFA, respectively). Plate #8 contains 200 μl of 1% NH4OH for elution. The beads are washed in loading buffer for 5 minutes at medium mixing speed, followed by binding of the phosphopeptides for 20 minutes and medium speed. The sequential washes are performed in 2 minutes and fast speed. Phosphopeptides are eluted in 10 minutes at medium mixing speed. After acidification, phosphopeptides were loaded directly on C18 evotips (Evosep) for MS analysis.

### Liquid chromatography-tandem mass spectrometry (LC-MS/MS) analysis

Label-free proteome and phosphoproteome samples were analyzed on the Evosep One system (Bache *et al*., 2018) coupled to an Orbitrap Exploris 480 (Bekker-Jensen, Martínez-Val, *et al*., 2020). Samples were separated on an in-house packed 15 cm analytical column (150 μm inner diameter), packed with 1.9 μm C18 beads, and column temperature was maintained at 60 ℃ using an integrated column oven (PRSO-V1, Sonation GmbH). Pre-programmed gradients were used: 30 samples per day for proteome, 60 samples per day for phosphoproteome. The mass spectrometer was operated in positive ion mode, using data-independent acquisition (DIA), as previously described (Bekker-Jensen, Bernhardt, *et al*., 2020), with spray voltage at 2 kV, heated capillary temperature at 275 °C and funnel RF frequency at 40. Full MS resolution was set to 120,000 at m/z 200 and full MS AGC target was 300%, with an injection time of 45 ms, and scan range was set to 350–1400 m/z. AGC target value for fragment scan was set at 1000%. 49 windows of 13.7 Da were used with an overlap of one Da. The MS/MS acquisition was set to 15,000 resolution, and injection time to 22 ms. Normalized collision energy was set at 27%. Peptide match was set to off, and isotope exclusion was on.

### Mass spectrometry raw data processing

Data were analyzed on Spectronaut V.15 in directDIA mode (spectral library-free) with the standard settings. For phosphoproteome analysis, the PTM localization filter was set at 0.75. Deamidation of asparagine and glutamine (NQ) was added as variable modification for both proteome and phosphoproteome data, and phosphorylation of serine, threonine and tyrosine (STY) only for the phosphoproteome data. The Human Uniprot fasta file (downloaded in 2019, 21,074 entries) was supplemented with a contaminant fasta files containing 246 entries.

### Bioinformatic analysis

Data analysis was done the Perseus Software (Tyanova *et al*., 2016) v1.6.15.0 and the Prostar online tool (http://www.prostar-proteomics.org/)(Wieczorek *et al*., 2017). Plots in Figs X and SX were performed using the ggplot2 package v3.3.5, using the R software package v4.1.1 (with RStudio v1.2.5042).

#### Proteome data

Protein group MS intensities were Log2-transformed and filtered by removing potential contaminants, rows without gene name and with less than 4 valid values in at least one condition (Ctrl, 1h or 4h). Data normalization was performed by variance stabilization normalization (VSN) as implemented in Prostar. Missing values were imputing by using the R plugin imputeLCMD (https://cran.rstudio.com/web/packages/imputeLCMD/index.html) as implemented in Perseus (QRILC=1). Inter-patient batch effect was removed by using the R plugin Combat(Johnson, Li and Rabinovic, 2007) as implemented in Perseus. Principal component analysis (PCA) was performed in Perseus on normalized, imputed and batch-corrected data. Differential expression analysis was performed in Perseus by using one-way ANOVA. p values were corrected by using the Benjamini–Hochberg (BH) procedure. Proteins were considered significantly regulated if p adjusted was < 0.01. Gene ontology (GO) and pathway (KEGG and Reactome) enrichment analyses were performed in Perseus by Fisher exact test against the whole dataset as background. p values were corrected by using the BH procedure. Hierarchical clustering of protein group intensities was performed in Perseus after row scaling (Z-score) by using Pearson correlation.

#### Phosphoproteome data

Phosphosite MS intensities were Log2-transformed and filtered by removing potential contaminants and phosphorylation sites with less than 4 valid values in at least one condition (Ctrl, 1h or 4h). The “expand the site table” function, implemented in Perseus, was used. MS intensities were normalized in Spectronaut. Missing value imputation (QRILC=1.3), batch effect removal, one-way ANOVA, GO/pathway enrichment analysis and hierarchical clustering were performed as described above for the proteome. Phosphosites were filtered based on their functional score with a cutoff of 0.45(Franciosa, Martinez-Val and Olsen, 2020). Functional protein network analysis was performed in STRING (Szklarczyk *et al*., 2019) through the STRING App (Doncheva *et al*., 2019) implemented for the Cytoscape software (Shannon *et al*., 2003). Network analysis (calculation of degree and closeness centrality) was performed in Cytoscape.

### Molecular Modelling: General Simulation Details

All simulations were performed with GROMACS 2019.3 (Abraham *et al*., 2015) at a 0.18 M NaCl salt concentration. Systems were kept at a constant temperature using a velocity rescaling thermostat (Bussi, Donadio and Parrinello, 2007) (τ_T_ = 1 ps). Pressure coupling was performed with the Berendsen barostat (Berendsen *et al*., 1998)(τ_T_ = 2 ps, 4.5·10^−5^compressibility and 1 bar reference pressure).

### Molecular Modelling: All-Atom Replica Exchange MD

All-atom molecular dynamics simulations were performed with the February 2021 version of the CHARMM36 force field (Huang and Mackerell, 2013) (Huang *et al*., 2016) with a 2 fs time step. The cofilin-1 NMR structure was downloaded from the PDB (entry 1tvj) and phosphorylated at Ser3 and Ser41 with the Charmm GUI(Lee *et al*., 2016). Protein structures were solvated in tip3p water and the systems were neutralized with additional Na^+^ or Cl^−^ ions. After the steepest descent minimization, the systems were equilibrated for 10 ns at 48 different temperatures, ranging from 310 to 427.5 K, with steps of 2.5 K. With these 48 equilibrated systems, replica-exchange runs were performed for 50 ns, with an exchange attempt every 500 steps. Van der Waals and Coulomb interactions were calculated using the shifted Verlet (Grubmüller *et al*., 2006) and particle mesh Ewald (PME) (Darden, York and Pedersen, 1998) methods, respectively, both with a 1.2 nm cut-off distance. The LINCS algorithm (Hess *et al*., 1997) was used to constraint bonds with hydrogen atoms. For analysis, the trajectories were de-multiplexed using GROMACS’ demux.pl script to obtain trajectories with continuous coordinates.

### Time-Lapse Microscopy

Day 6 moDCs were transfected with GFP-WT cofilin or mCherry-labelled mutant cofilin using a Neon electroporator (Thermofisher Scientific). moDCs were pulsed twice at 1000 V for 40 ms and using 10 µg of the corresponding plasmid. After transfection cells were immediately incubated at 37 °C for 4 h in phenol red-free RPMI 1640 containing 10 % FBS and 1 % ultra-glutamine prior to being sorted using a FACS Melody WT or mutant cofilin expressing cells were then embedded into a collagen matrix with a final concentration of 1.7 mg ml^−1^. Collagen matrix was generated using a PureCol® Type I Bovine Collagen Solution (Advanced Biomatrix), α-modified minimal essential medium (Sigma Aldrich) and sodium bicarbonate. Collagen mixture was allowed to pre-polymerise for 5 minutes at 37 °C prior to adding a 45 µl cell suspension containing 60.000 moDCs expressing either WT or mutant cofilin in phenol red–free RPMI. The total mixture was transferred to a 96-well black plate (Greiner Bio-One; 655090) and incubated for 45 minutes at 37°C. Afterwards, phenol red–free RPMI 1640 culture medium supplemented with 1 % ultraglutamine and 10 % FBS was added on top of the matrices.

Time lapsed video microscopy was performed using the Zeiss Axio Observer 7 with Zeiss Axiocam 702 camera, Sutter Lambda DG5 light source, fast filter wheels, Zen image acquisition and analysis software and Ibidi stage incubator. Sequential images were taken every 5 min using an LD Plan-Neofluar 20x /0.4 Korr M27 GFP_WT cofilin expressing cells were excited with a 488 laser and mCherry_mutant cofilin cells with a 592 laser.

Individual cell tracking was performed to determine the median cell velocity, the cell mean square displacement (MSD) and the Euclidean distance reached by a cell after 120 minutes of tracking. Cell tracking was performed 4 hours after embedding cells in the collagen to allow adaptation to the environment and using the manual tracking plugin of Fiji (ImageJ) with adjusted microscope-specific time and calibration parameters. Cells were chosen randomly and dead cells were excluded from the cell tracking. The MSD over time intervals was determined as described in the protocol of Van Rijn et al. (van Rijn et al. 2016). In short, the MSD was calculated per time interval for each cell. The average per time interval was calculated for all cells corrected for the tracking length of the cells. The Euclidean distance was calculated using the Chemotaxis and Migration software (Ibidi).

## Data availability

All raw MS data were generated by the authors and deposited to the ProteomeXchange Consortium (http://proteomecentral.proteomexchange.org) via the PRIDE partner repository (Perez-Riverol *et al*., 2022) with the dataset identifier PXD031056. Primary microscopy data generated in this study have been deposited in the Zenodo database under accession code 10.5281/zenodo.7081102.

## Acknowledgments

We are thankful to P. Grijpstra for technical support throughout the project. We are also grateful to F. Stempels for advice regarding FLIM observations. G.v.d.Bogaart is supported by the European Research Council (ERC) under the European Union’s Horizon 2020 research innovation program [grant agreement No. 862137]; and ZonMW [project grant No. 09120011910001]. The proteomics experiments performed in this project and GF’s salary were funded by the European Union’s Horizon 2020 research and innovation program under grant agreement EPIC-XS-823839. Work at the NNF CPR is funded by a donation from the NNF (NNF14CC0001). CK is funded by the European Union’s Horizon 2020 research and innovation program under the Marie Sklodowska-Curie grant agreement No.861389. L.Q.Cano is supported by grant 11618 from the Dutch Cancer Society. WHR is supported by a FOM projectruimte grant.

---

**Figure S1.**
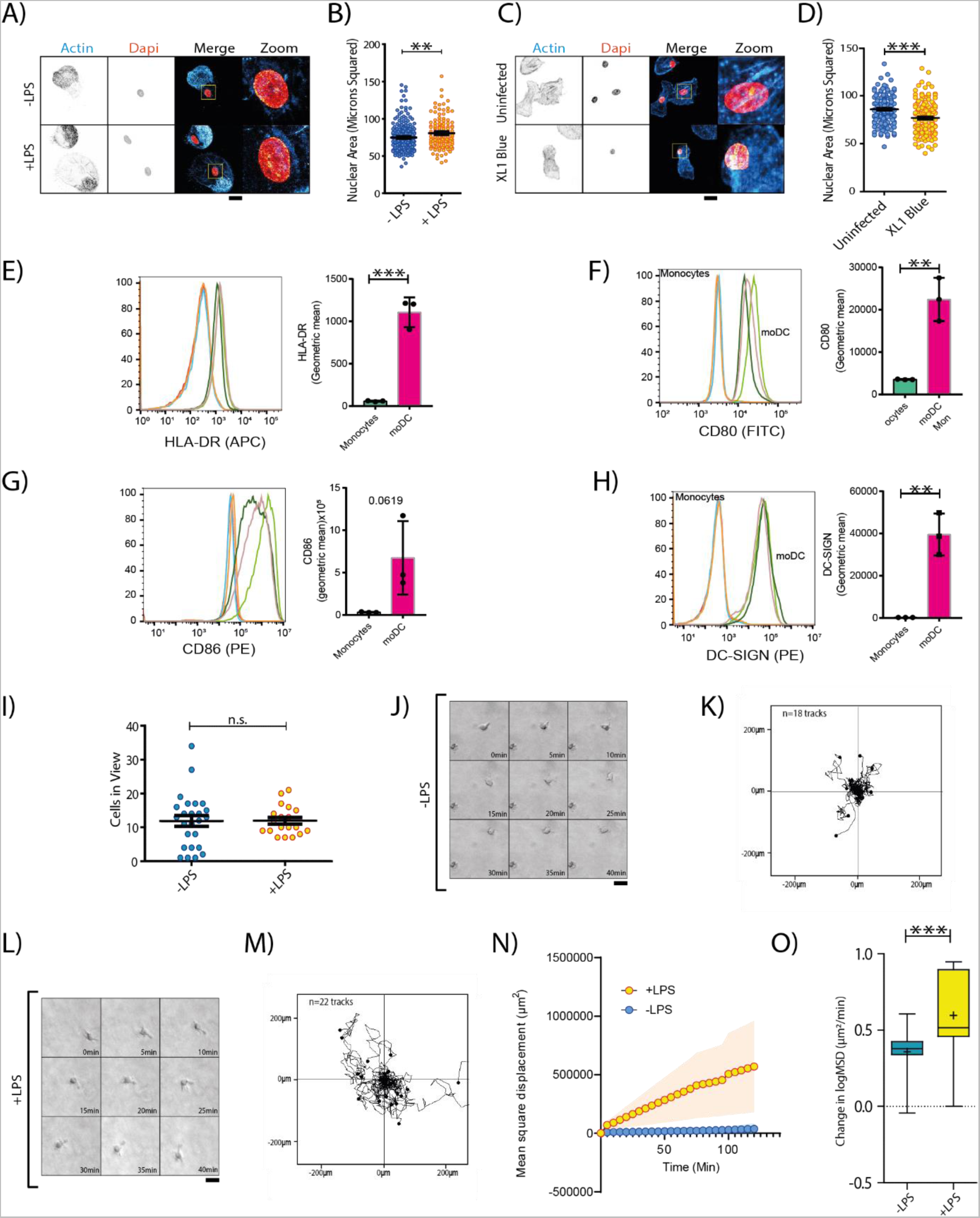
Inflammation Driven Nuclear Re-Programming is Dendritic Cell Specific. A) Confocal micrographs of CD14^+^ macrophages stained with phalloidin (cyan in merge) and DAPI (orange), and cultured overnight in the presence or absence of LPS. B) Quantification of the Z-projected nuclear areas of CD14^+^ macrophages (minimum of 122 measurements per condition over 3 biological repeats). C) Confocal micrographs of dendritic cells cultured overnight in the presence or absence of E.coli. D) Quantification of Z-projected nuclear area. (minimum of 174 measurements over 3 biological repeats). E) Quantification of DC-SIGN expression for monocytes and moDCs reveals that DC-SIGN expressing cells are largely absent in monocyte populations prior to differentiation. Curves on flow cytometry histogram (left) represent separate donors. F) Quantification of HLA-DR expression for monocytes and moDCs reveals that HLA-DR expressing cells are largely absent in monocyte populations prior to differentiation. Curves on flow cytometry histogram (left) represent separate donors. G) Quantification of CD80 expression for monocytes and moDCs reveals that CD80 expressing cells are largely absent in monocyte populations prior to differentiation. Curves on flow cytometry histogram (left) represent separate donors. H) Quantification of CD86 expression for monocytes and moDCs reveals that CD86 expressing cells, whilst present in monocyte populations increase (non-significantly – p=0.0619) following differentiation. Curves on flow cytometry histogram (left) represent separate donors. I) Dendritic cells in view in set imaging field size in the absence or presence of LPS (data from at least 19 fields of view across 3 donors). J) Example movie of inactive dendritic cell migrating in collagen. K) Examples migration traces of inactive dendritic cells. L) Example movie of active dendritic cell migrating in collagen. M) Examples migration traces of active dendritic cells. N) Example MSDs of inactive (-LPS) and active (+LPS) dendritic cells migrating in collagen. Shaded region represents SEM. O) Quantification of rate of MSD change per donor, for inactive and active dendritic cells (90 measurements per condition over 3 biological repeats). Data information: Scale bars indicate 20 microns unless otherwise stated. Statistical significance calculated using 2-sided unpaired t-tests for 2 condition experiments or ANOVA/Tukey multiple comparison tests for 3+ condition experiments (with test selected according to distribution pattern of the data). **, P < 0.01; ***, P < 0.001. Error Bars= SEM. For Box and Whisker plots, Box represents 25th to 75th percentiles, whiskers represents maximum and minumum values, middle band represents data median,+ represents data mean.

**Figure S2.**
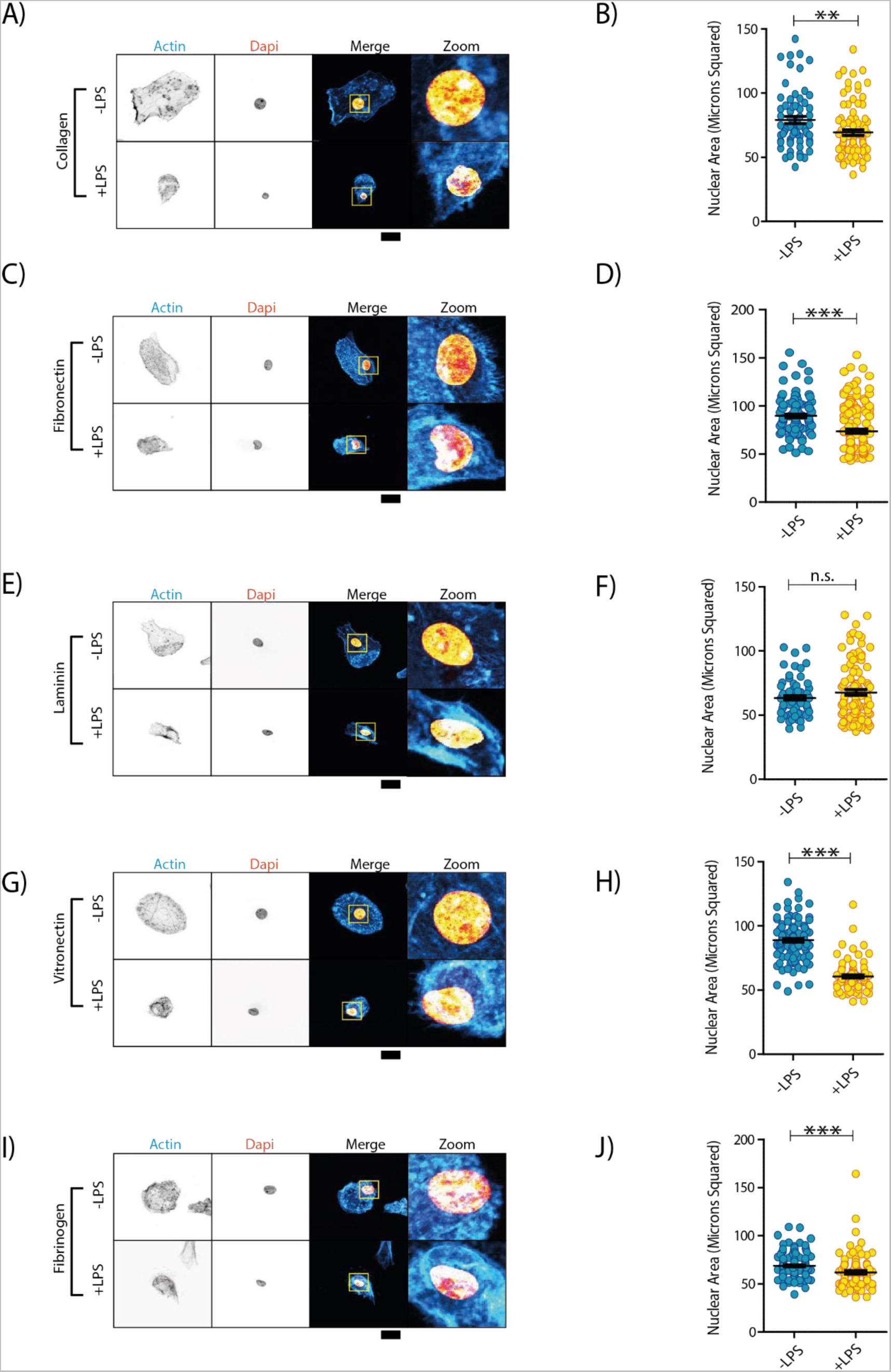
Spherical Deformation of the Nucleus is Typically Unaltered by ECM Proteins. A) Confocal micrographs of dendritic cells stained with phalloidin (cyan in merge) and DAPI (orange), and cultured in the presence or absence of LPS (overnight stimulation) on collagen-coated glass. B) Quantification of Z-projected nuclear areas of dendritic cells cultured on collagen-coated glass (minimum of 67 measurements per sample over 3 biological repeats). C) Confocal micrograph of dendritic cells cultured in the presence or absence of LPS on fibronectin-coated glass. D) Quantification of Z-projected nuclear areas of dendritic cells cultured on fibronectin-coated glass (minimum of 54 measurements per condition over 3 biological repeats). E) Confocal micrograph of dendritic cells cultured in the presence or absence of LPS on laminin-coated glass. F) Quantification of Z-projected nuclear areas of dendritic cells cultured on laminin-coated glass (minimum of 47 measurements per condition across 3 biological repeats). G) Confocal micrograph of dendritic cells cultured in the presence or absence of LPS on vitronectin-coated glass. H) Quantification of Z-projected nuclear area of dendritic cells cultured on vitronectin-coated glass (minimum of 81 measurements per condition over 3 biological repeats). I) Confocal micrograph of dendritic cells cultured in the presence or absence of LPS on fibrinogen-coated glass. J) Quantification of Z-projected nuclear areas of dendritic cells cultured on fibrinogen-coated glass (minimum of 104 measurements per condition over 3 biological repeats). Data information. Scale bars indicate 10 microns. Scale bars indicate 20 microns. Statistical significance calculated using 2-sided unpaired t-tests (with test selected according to distribution pattern of the data). **, P < 0.01; ***, P < 0.001; n.s., not significant. Error Bars=SEM.

**Figure S3.**
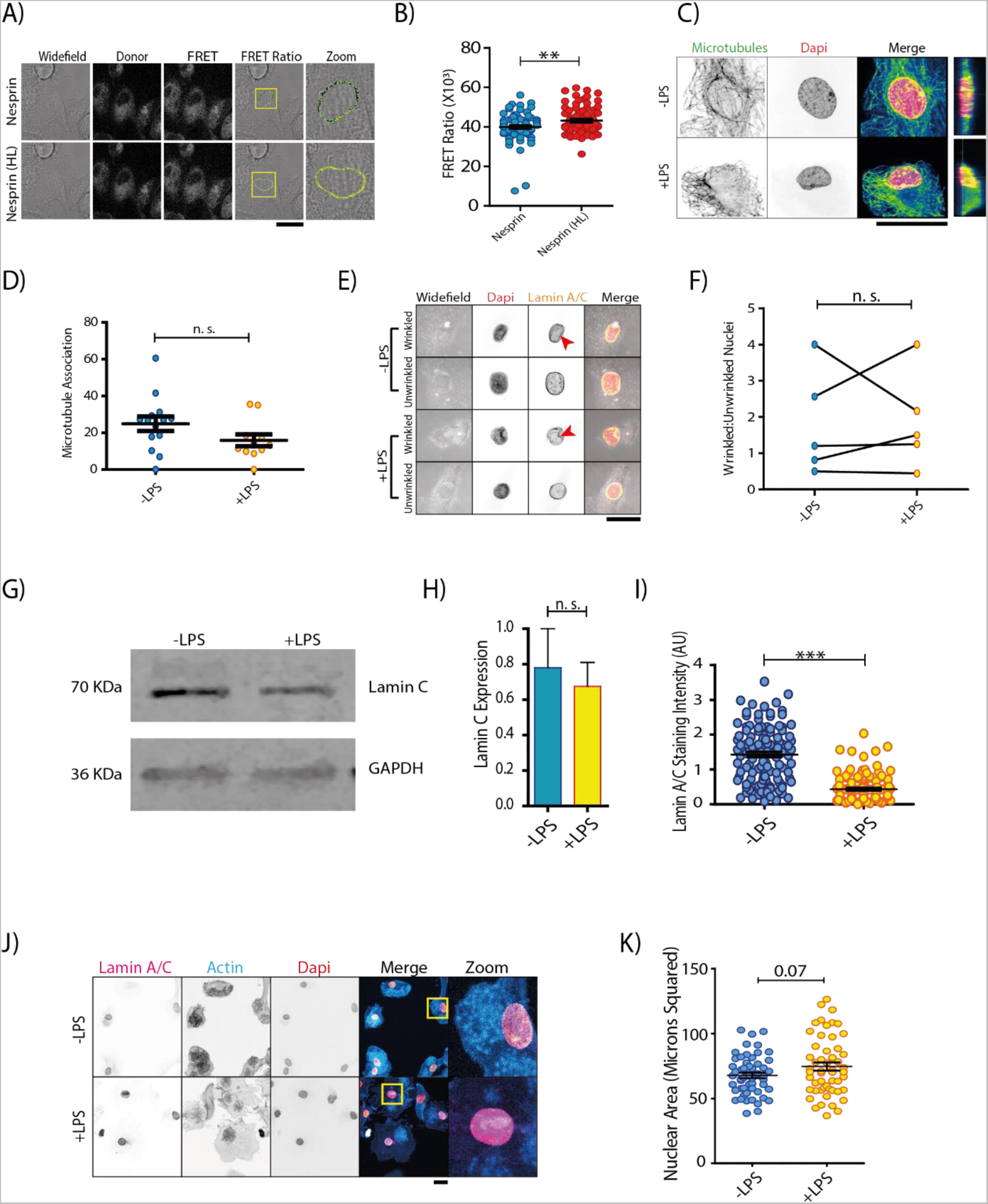
A) Example ratiometric FRET images of Nesprin-2 WT and headless probes, expressed in dendritic cells. B) Quantification of FRET ratio at the dendritic cell nuclear membrane (minimum of 79 measurements per condition over 3 donors). C) Airyscan images of immunostained microtubules (green in merge) around the dendritic cell nucleus in the absence or presence of LPS. Cyan: phalloidin. Organe: DAPI. D) Quantification of average microtubule intensity above dendritic cell nuclei (at least 11 measurements per condition over 4 donors). E). Immunofluorescence images of lamin A/C (yellow in merge) in dendritic cells, showing wrinkled and non-wrinkled nuclei in the presence or absence of LPS (overnight stimulation). F) Quantification of the change of ratio of Lamin-A/C wrinkled to unwrinkled dendritic cell nuclei, following LPS stimulation (overnight) (data from 5 donors). G) Western blot of Lamin-A/C. Dendritic cells primarily expressed lamin-C. GAPDH: loading control. H) Quantification of Lamin C levels using western blot (normalized to GAPDH) in dendritic cells reveals a small and non-significant change in reduction Lamin-C expression as a result of dendritic cell activation (data from 4 donors). I) Quantification of lamin-A/C intensity from confocal micrographs (set laser power and detector sensitivity), reveals that activation reduces lamin-A/C levels within dendritic cell nuclei. J) Immunofluorescence images of dendritic cells overexpressing lamin-A/C in the absence or presence of LPS (minimum of 122 measurements per condition over 3 donors). K) Quantification of nuclear size reveals that overexpression of lamin-A/C is sufficient to block the nuclear deformation phenotype associated with dendritic cell activation (minimum of 53 measurements per condition over 3 donors). Data information Scale bars indicate 20 microns. Statistical significance calculated using an unpaired t-test for 2 condition experiments or ANOVA/Tukey multiple comparison test for 3+ condition experiments **, P < 0.01; ***, P < 0.001; n.s., not significant.

**Figure S4.**
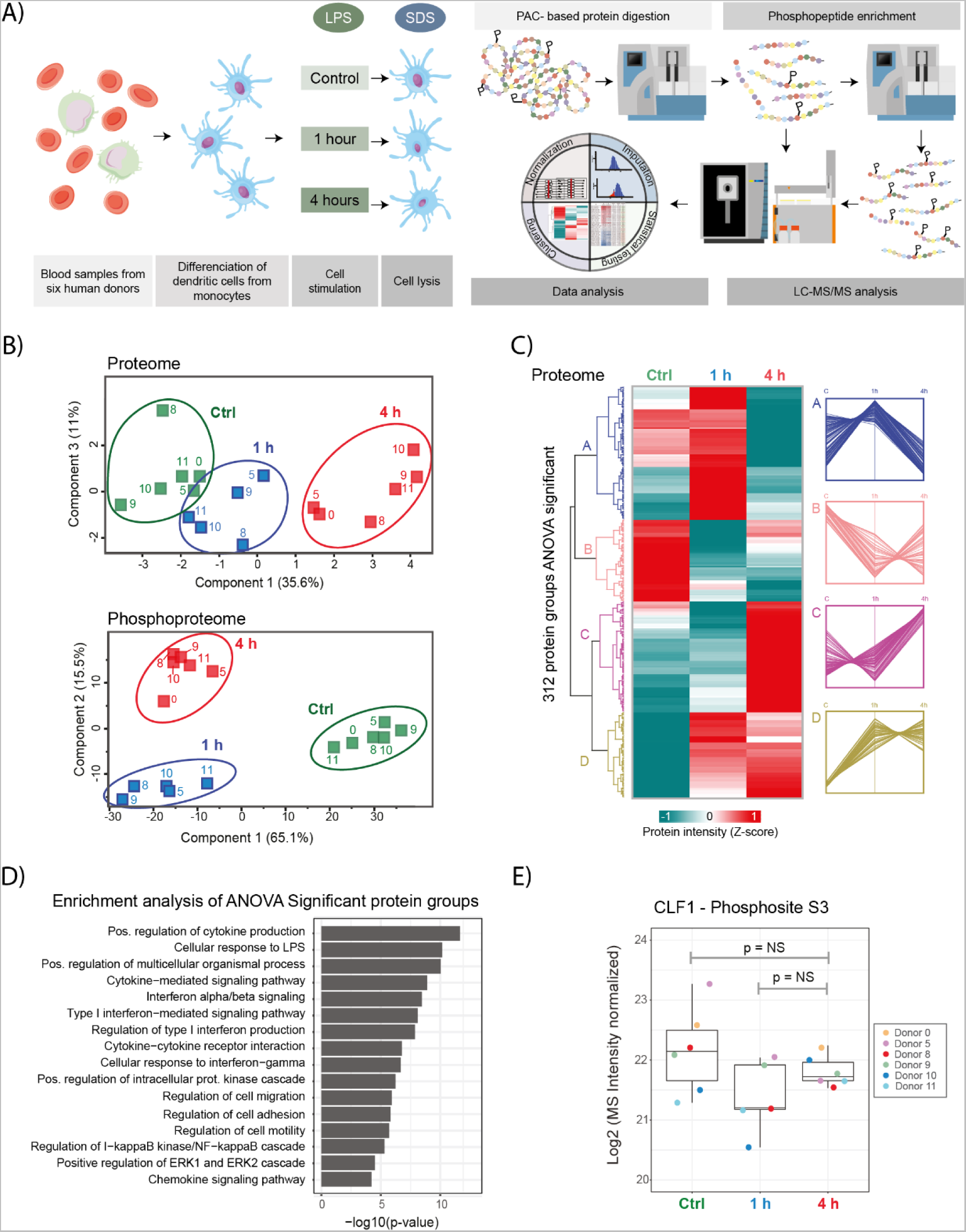
Dendritic Cell Proteomics Outline. A) Depiction of the phosphoproteomics workflow. B) Principal component analysis on normalized, imputed and batch-corrected MS signal. Proteome: up. Phosphoproteome: down. C) Hierarchical clustering analysis by using Pearson correlation distance of protein intensities. Values were normalized, imputed, batch-corrected and scaled (Z-score) before clustering. Only sites with one-way ANOVA FDR < 0.01 were used for clustering. D) Gene ontology and Pathway (KEGG and Reactome) enrichment analysis on all proteins shown in C. Enrichment was performed against the entire proteome quantified. E) Boxplot of the normalized MS signal, after Log2 transformation, for the phosphorylated serine 3 of cofilin-1. Data information: P values were calculated by two-sided paired Student’s t test. NS: not significant. For Box and Whisker plots, Box represents 25th to 75th percentiles, whiskers represents maximum and minumum values, middle band represents data median.

**Figure S5.**
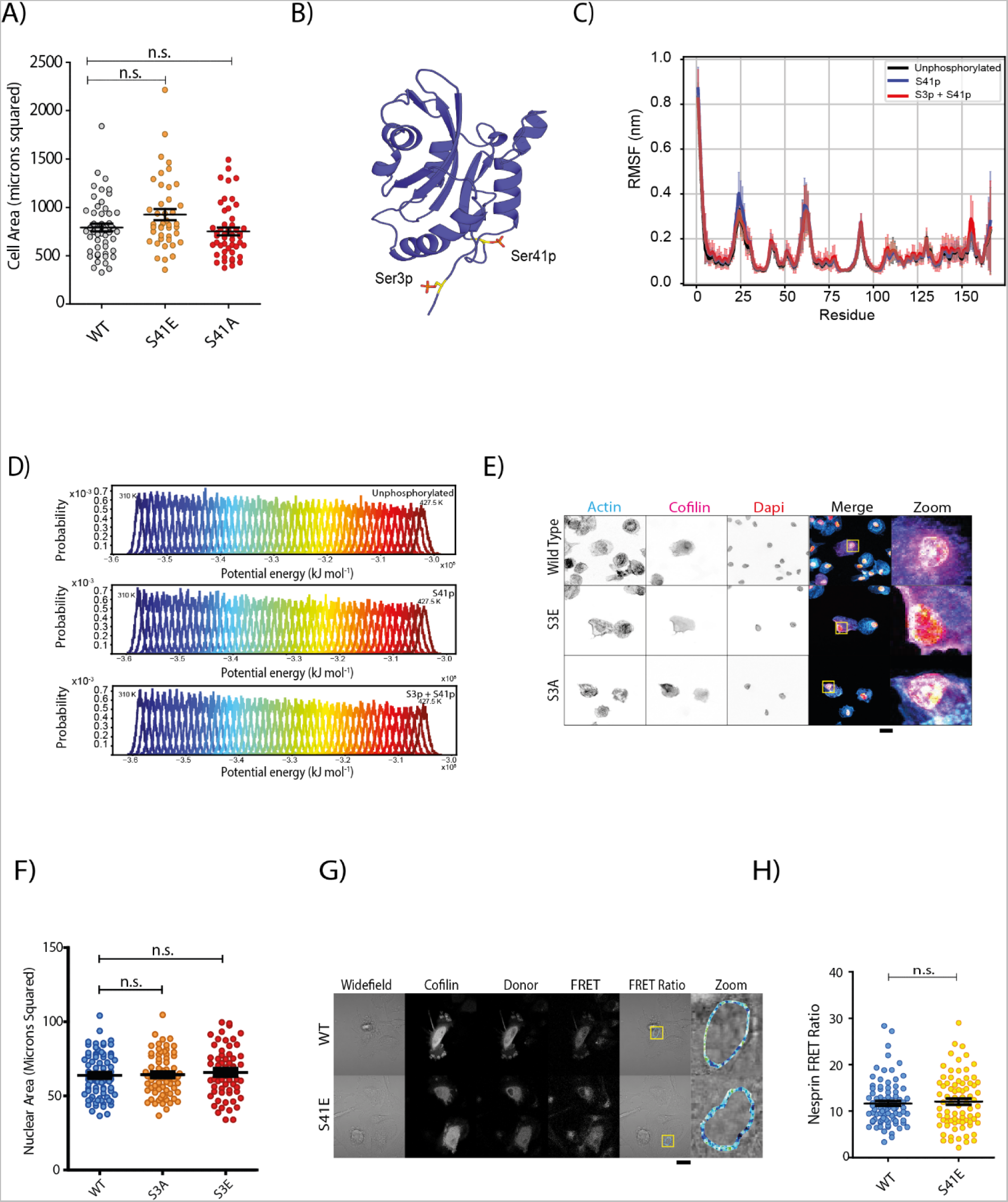
Cofilin-Mediated Nuclear Deformation is Independent of Altered Cell-Spreading or LINC Complex Reprogramming. A) Total Z-projected area of dendritic cells expressing WT or S41E and S41A cofilin-1 mutants (minimum of 38 measurements per condition over 3 biological repeats). B) NMR-structure of cofilin-1, with sites S3 and S41 highlighted. RCSB protein databank structure 1TVJ (Gorbatyuk et al., 2006) C) Temperature replica exchange molecular dynamics (REMD) simulations examining the effects of S3 and S41 phosphorylation on the dynamics of cofilin-1 reveals that no combination of phosphorylation/de-phosphorylation events (of S3 and S41) alters the core structure of cofilin-1. D) Energy distributions in the temperature range (310-427.5 K) used for REMD simulations. E) Confocal micrographs of dendritic cells expressing GFP-tagged WT, S3E or S3A cofilin-1 (magenta in merge). Blue: phalloidin. Orange: DAPI. F) Quantification of the Z-projected nuclear areas revealed that neither the S3E (phoshomimetic) nor S3A (phosphodead) variants can induce deformation of the dendritic cell nucleus (minimum of 60 measurements per condition over 3 biological repeats). G) Example ratiometric FRET image of Nesprin-2 probe, expressing either WT or S41E cofilin. H) Quantification of FRET ratio at the dendritic cell nuclear membrane reveals that S41E cofilin mediated nuclear remodelling occurs independently of the removal of actin-based LINC complexes across of the dendritic cell nuclear membrane (minimum of 77 measurements per condition over 3 biological repeats). Data information: Scale bars indicate 20 microns. Statistical significance calculated using an Statistical significance calculated using an unpaired t-test for 2 condition experiments or ANOVA/Tukey multiple comparison test for 3+ condition experiments (with test selected according to distribution pattern of the data). **, P < 0.01; ***, P < 0.001; n.s., not significant. Error Bars= SEM.

